# Increased medial collagen enhances aortic resilience against mural delamination from hydraulic fracturing

**DOI:** 10.64898/2026.05.12.724717

**Authors:** Alan Chou, Kristina Wang, Dustin Lieu, Prashanth Vallabhajosyula, Jay D. Humphrey, George Tellides, Roland Assi

## Abstract

The aorta, normally resilient to hemodynamic stresses, becomes vulnerable to structural failure due to diverse conditions that weaken the wall. We injected fluid into excised specimens of human ascending aorta with pressure monitoring to quantify the impact of clinical and histological factors on mural damage. Two modes of medial injury emerged with distinct pressure tracings. Extravasation was characterized by diffuse infiltration of fluid with widespread damage of smooth muscle cells and collagen fibers but limited separation of elastic lamellae. By contrast, delamination was characterized by marked separation of elastic lamellae along a single plane with damage to cells and fibrillar matrix restricted to adjacent laminae. Aging, aortic dilatation, and family history associated with lower pressures causing delamination, whereas a diagnosis of hypertension associated with higher pressures suggesting resilience to dissection. Collagen fraction adjacent to delamination correlated with higher pressures as did decreased smooth muscle cell density and increased glycosaminoglycan fraction, although several clinical and histological variables were interrelated. Protein cross-linking strengthened and enzymatic digestion of collagen weakened the aortic wall, while acute cell lysis with detergent had no effect. We conclude that increased functional medial collagen has an adaptive protective role in aortic remodeling rather than signifying medial degeneration.

## Introduction

Ascending aortic dissection is a catastrophic event in which tears of the aortic wall allow entry of blood between the layers of the media that frequently results in interrupted blood flow to vital organs or rupture of the aortic wall. If afflicted individuals survive until hospitalization, lifesaving treatment is emergent, high-risk surgical replacement of the dissected proximal aorta with a synthetic conduit (1, 2). Consequently, prophylactic replacement of vulnerable aortas is recommended prior to loss of structural integrity. Current guidelines recognize maximal aortic diameter and rapid growth as primary indications for intervention with lower thresholds for patients with heritable connective tissue disorders, such as Marfan or Loeys-Dietz syndromes (3). Other associated risk factors such as advanced age, hypertension, smoking, and bicuspid aortic valve are not included in guidelines for elective repair as the associated increase in dissection risk is unclear (4–6). Despite the complexity of risk prediction, guidelines remain largely based on retrospective studies measuring aortic diameter at the time of dissection (7, 8). Nonetheless, approximately half of patients with ascending aortic dissections have aortic diameters below the threshold for surgical replacement (9, 10). Thus, there is considerable need for improved methods to identify aortas vulnerable to dissection.

Smooth muscle cells and extracellular matrix – including elastic lamellae, fibrillar collagens, and glycosaminoglycans – govern functional and structural properties of the aortic media (11, 12). Investigators have developed biomechanical tests of aortic tissue to quantify resistance to structural failure (13, 14). Planar biaxial tests and bulge inflation tests impose elongation or pressurization of a segment of aorta under well-controlled conditions with automated systems. These tests are limited, however, in their ability to predict aortic dissection as their typical method of failure is transmural rupture. Shearing and peeling tests involve forced separation of adjacent laminae of the media, termed delamination, but impose non-physiologic conditions on the tissue by restricting the direction of damage. A classical test introduced in the 19th century, reintroduced in the mid-20th century with pressure monitoring, and further studied by Margot Roach in the 1990s forcibly delaminates the aortic media by intramural injection of fluid (15–19). This test allows dissection to be simulated via a clinically relevant mechanism of hydraulic fracturing in intact aortic tissue. Among other findings, prior investigators demonstrated that supraphysiologic pressures are required to initiate the dissection but pressures required for propagation of the dissection are commonly within the physiological range.

The present study was motivated by the need for a quantitative method to assess clinical and structural factors that determine vulnerability of human aortas to dissection. We confirm that ex vivo intramural fluid injection is a simple, reproducible method for medial delamination that mimics critical aspects of in vivo aortic dissection. We also describe an unappreciated manifestation of aortic wall damage from hydraulic fracturing termed medial extravasation. By testing aortic specimens from a broad range of subjects, we examine correlations of clinical factors and histological features with pressures required for delamination. Surprisingly, a diagnosis of hypertension and certain features of medial degeneration correlate with resilience against, not vulnerability to, dissection. Mechanistic experiments confirm that greater collagen cross-linking and content, but not acute loss of smooth muscle cells, increase delamination pressures. We interpret medial fibrosis as a reparative response of smooth muscle cells to stress and injury rather than signifying structural deterioration.

## Results

### Hydraulic fracturing of the aortic wall results in medial extravasation or delamination

To examine propensity to aortic dissection, we injected fluid (normal saline with 40 g/mL human albumin and 1% India ink v/v) into ascending aorta specimens from organ donors and patients undergoing elective surgery for thoracic aortic aneurysms. Based on pilot data from 5–10 specimens in which the type of medial injury was similar at flow rates of 30 and 100 μL/min but unlike those of 300 and 600 μL/min, we chose 100 and 300 μL/min as representative of low and high injection rates with different outcomes. Simultaneous pressure and video monitoring revealed two distinct types of medial damage: extravasation and delamination. Medial extravasation was characterized by expansion of the dye-colored area at a constant rate with hazy, irregular outlines (Figure 1A). Fluid spread in axial and circumferential directions with minor thickening of the aortic wall. The injection pressure rose steadily until reaching a plateau with a slight upslope, consistent with the constant rate of fluid accumulation (Figure 1B). Histology showed minor (< 100 μm) separation of elastic lamellae extending from central to distant zones of dye infiltration, ranging from focal detachments near the site of injection to areas with mild-to-moderate separation interspersed with areas without separation of lamellar structures (Figure 1C). By contrast, medial delamination was characterized by minimal dye entry during early injection, followed by the sudden appearance of a bleb that subsequently expanded at a constant rate in axial, circumferential, and radial directions (Figure 1D). There was marked thickening of the aortic wall due to pooling of fluid within the media. The pressure history showed a sharp initial rise to a high peak, followed by a rapid drop corresponding to initial bleb formation, then a final plateau phase with slight downslope corresponding to further bleb expansion (Figure 1E). Histology showed a single plane of widely separated elastic lamellae considered as major delamination (up to several mm) with minimal staining by dye beyond the cavity edge (Figure 1F). Outside of the first few elastic laminae bordering the delamination plane, the inner and outer media adjacent to the damaged area appeared intact. Most injections yielded patterns of damage that matched either extravasation or delamination in all three categories (gross appearance, pressure tracing, and histological features), though intermediate manifestations were seen in a few cases (∼3%) and were excluded from analysis.

**Figure 1:**
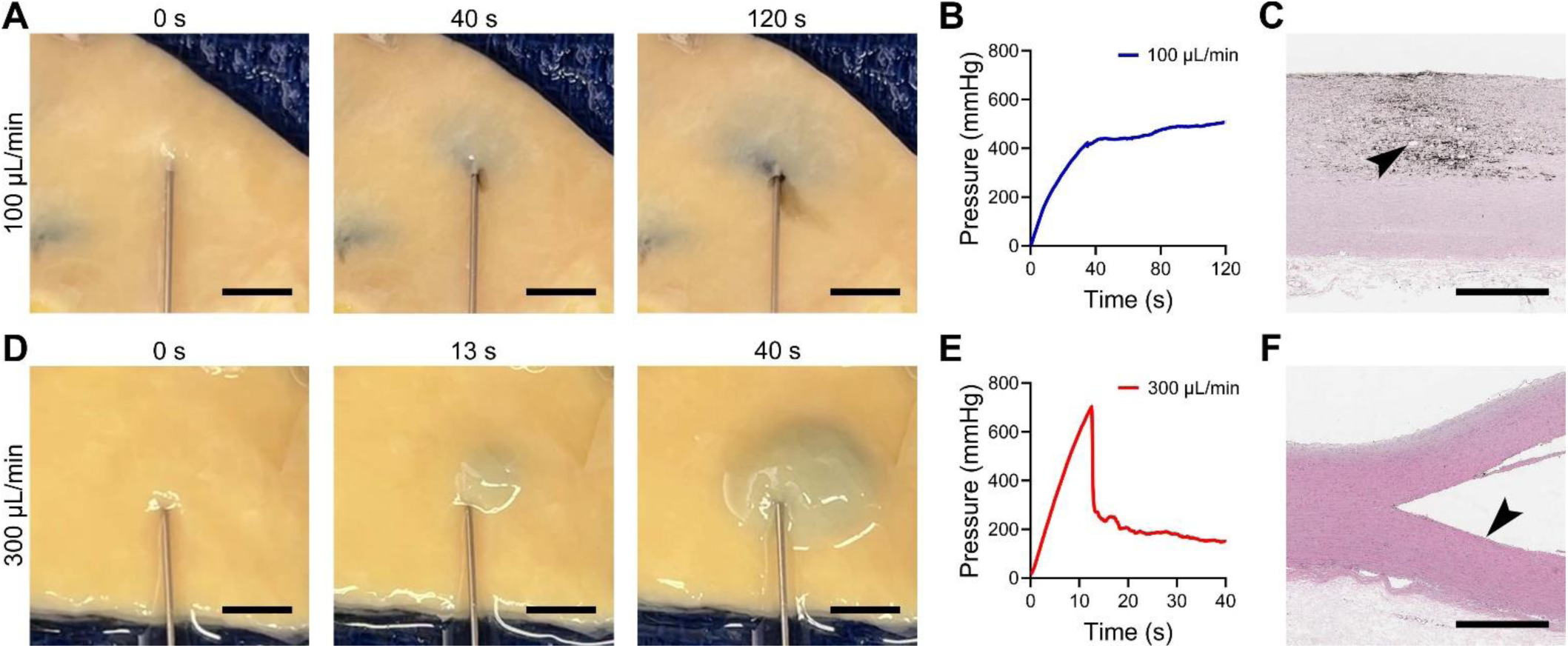
Medial extravasation or delamination after intramural fluid injection. Intramural injection of saline containing albumin and India ink into aortic specimens at different flow rates of 100 μL/min for 120 s or 300 μL/min for 40 s resulted in either medial extravasation (upper panels) or medial delamination (lower panels), respectively. For extravasation, (**A**) snapshots from video monitoring of the intimal surface showed steady expansion of the dye-discolored area with irregular borders but no bleb formation, (**B**) pressure-time history monitoring showed a steady rise in perfusion pressure to a plateau with minimal further increase corresponding to steady expansion of the extravasation area, and (**C**) histology with hematoxylin and eosin stain showed diffuse extravasation of India ink into the aortic media with few focal separations of laminae appearing as small defects (arrowhead) near the needle site. Conversely, delamination was characterized by (**D**) slow initial expansion of the dye-discolored area followed by rapid bleb expansion, (**E**) rapid initial increase in perfusion pressure reaching a peak, followed by a rapid decrease corresponding to bleb formation, and then a plateau with minimal further decrease corresponding to bleb enlargement, and (**F**) substantial separation of laminae along a single plane with minimal extravasation of India ink into the subjacent media (arrowhead). Scale bars represent 5 mm (panels A and D) or 1 mm (panels C and F). Data are representative of 102 aortic specimens injected with fluid.

### Extravasation and delamination cause distinct structural damage of the media

Confocal immunofluorescence microscopy revealed characteristics of tissue damage resulting from hydraulic fracturing of the media. In medial extravasation, we observed limited, or no, separation of elastic lamellae from the center to periphery of lesions (Figure 2A). This damage included areas with complete loss of smooth muscle cells and areas with residual attached cell fragments containing smooth muscle α-actin filaments but without nuclei. Most separated elastic lamellae had attached fragments of collagen fibers, but not any increase in elastin breaks. In medial delamination, fragmentation of cells and fibrillar matrix was restricted to the edge of the bleb and, to a lesser degree, within a few adjacent laminae (Figure 2B). Parts of the exposed elastic lamellae along the major delamination plane were stripped clean of smooth muscle cells and collagen fibers, with occasional elastin breaks found in adjacent lamellae. Cells and fibrillar matrix beyond the third lamina from the major delamination plane usually appeared intact. In summary, extravasation causes diffuse minor separation of elastic lamellae and widespread damage to medial cells and fibrillar matrix, while tissue injury from delamination is largely confined to the major delamination plane and directly adjacent laminae.

**Figure 2:**
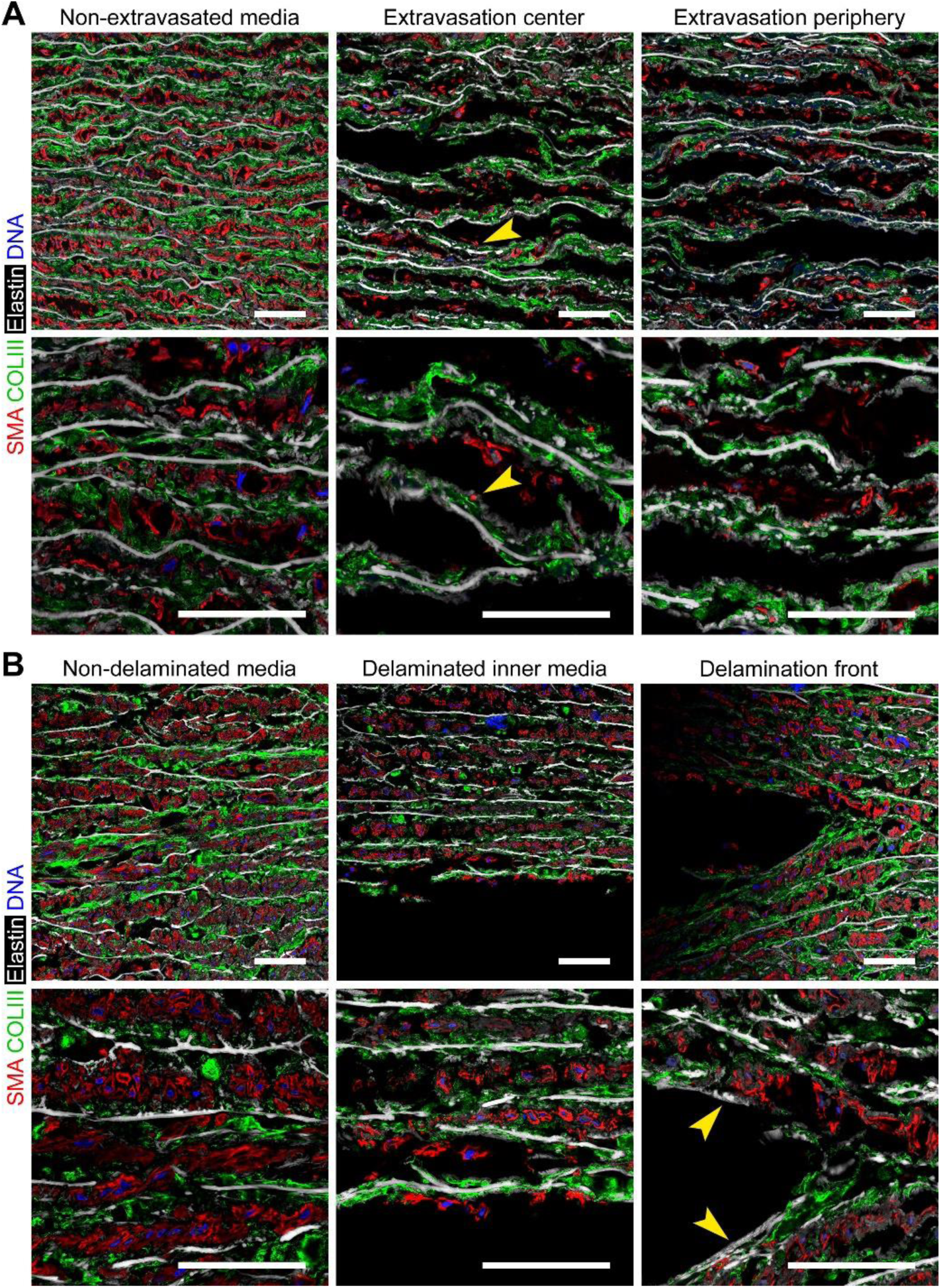
Structural damage from medial extravasation vs. delamination. Fluid-injected aortic specimens with extravasation or delamination were imaged by confocal immunofluorescence microscopy after labeling with antibodies to smooth muscle α-actin (SMA, red) and type III collagen (COLIII, green), Alexa Fluor 633 hydrazide for elastin (white), and DAPI for DNA (blue). (**A**) Extravasation results in mild-to-moderate separation of many, though not all, laminae from lesion center to periphery associated with displacement and fragmentation of smooth muscle cells and collagen fibers with numerous fragments remaining attached to elastic lamellae (arrowheads). (**B**) Delamination results in severe medial damage, including stripping of some elastic laminae free of cell and fibrillar matrix fragments (arrowheads), limited to the major delamination plane with mild delamination and moderate damage of cells and collagen fibers in the adjacent 2–3 laminae; the deeper subjacent media appeared intact. Scale bars represent 50 μm. Data are representative of 6 specimens examined by confocal microscopy.

### Medial extravasation or delamination depend on flow rate and subject

We examined for experimental and subject factors that associate with medial extravasation versus delamination. Subject characteristics are summarized in Supplemental Table 1 and the number of subjects, specimens, and injections are shown in Supplemental Figure 1. Surgical patients were more likely to have dilatated aortas, aortic valve abnormalities, and a family history of thoracic aortic aneurysm and dissection, while organ donors had higher rates of diabetes and smoking. Fresh aortic specimens from 65 subjects were tested by intramural fluid injection, including separate specimens from proximal and distal regions of the ascending aorta from a subgroup of subjects yielding 102 specimens from distinct axial locations with unique aortic diameters. Overall, 101 specimens were injected at high flow rate and 73 specimens were injected at low flow rate (including injections of 72 specimens at both 100 and 300 μL/min via separate sites, 29 specimens at 300 μL/min only, and 1 specimen at 100 μL/min only). Pressures from replicate injections at the same flow rate (range of 1–6 injections depending on the size of the specimen) were averaged and reported as single values. Extravasation was more frequent in aortas injected at low flow rate, while delamination predominated at high flow rate (Figure 3A). Aortas from younger subjects were predisposed to extravasation rather than delamination at both flow rates (Figure 3B). Additionally, dilated aortas were more likely to delaminate at high flow rate (Figure 3C). Six specimens that only extravasated at both high and low flow rates were from younger subjects (43.0 ± 21.1 years) with smaller aortas (33.5 ± 7.3 mm), while 30 specimens that only delaminated at both high and low flow rates were from older subjects (68.1 ± 8.8 years) with larger aortas (41.2 ± 5.5 mm). Thus, delamination as a mode of medial injury correlates with known clinical risk factors for aortic dissection, namely, aging and aortic dilatation, whereas medial damage from extravasation occurs in the absence of these risk factors.

**Figure 3:**
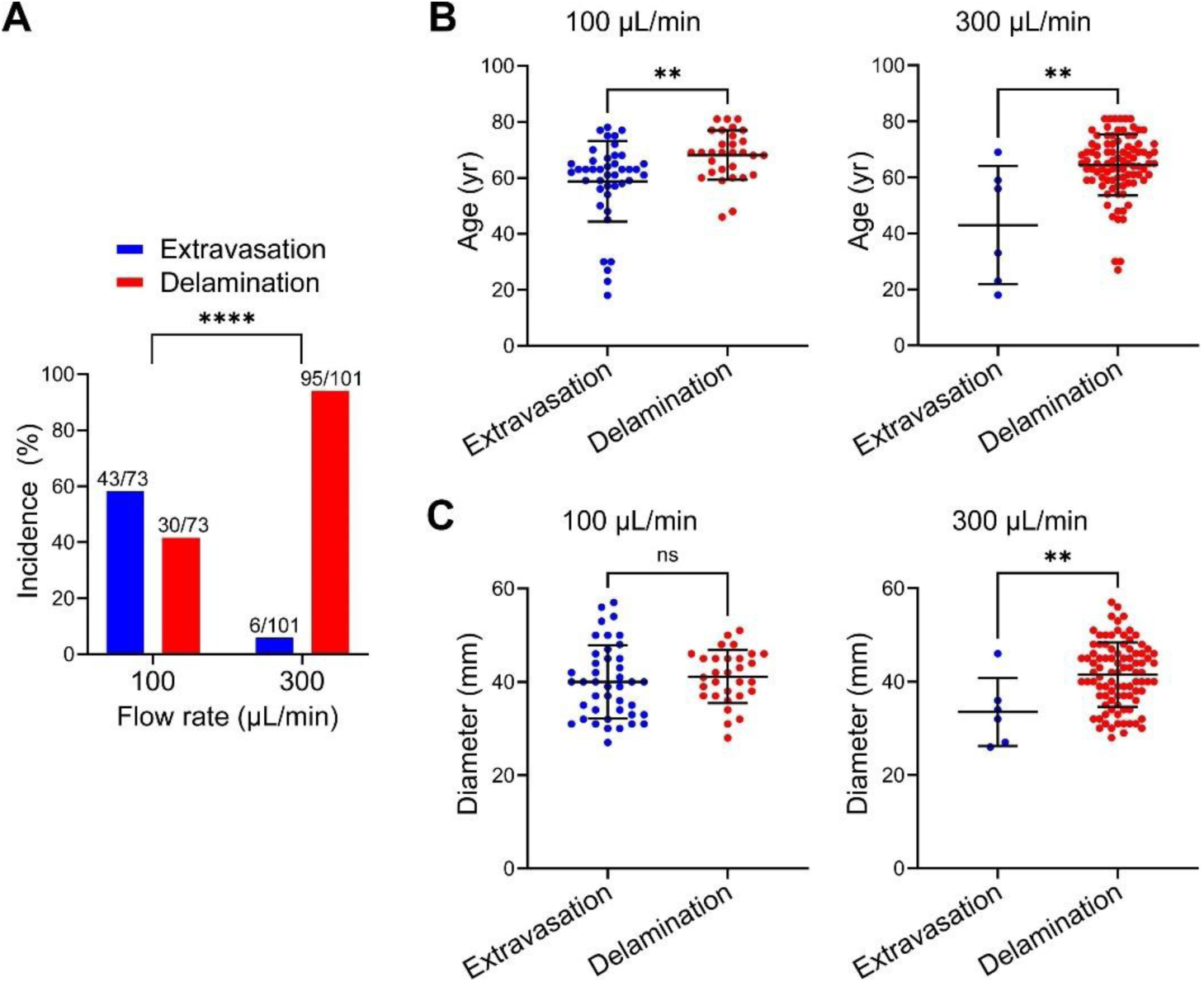
Determinants of medial extravasation vs. delamination. The type of medial injury depended on methodological and subject factors. (**A**) Extravasation was more common with intramural fluid injection at 100 μL/min, whereas delamination was more frequent at 300 µL/min. (**B**) Aortas from older subjects were more likely to delaminate than extravasate at both low and high flow rates. (**C**) Greater maximal ascending aortic diameter predisposed to delamination over extravasation at high but not low flow rates. Data for individual specimens are shown with lines representing mean ± SD; *n* = 73 at 100 μL/min, *n* = 101 at 300 μL/min, ns: not significant, \*\**p* < 0.01, \*\*\*\**p* < 0.0001, Fisher’s exact test (panel A), Mann-Whitney test (panel B), and unpaired t-test (panel C).

### Increased vulnerability of the greater curvature to medial delamination

We exploited quantitative features of injection pressure histories during delamination to assess possible regional susceptibility of the ascending aorta to hydraulic fracturing. Specimens injected at high flow rate with minimal leakage that resulted in well-defined delamination with a characteristic rapid increase, peak, and then decline to a plateau in pressure were included for analysis; a total of 216 injections in 90 specimens from 56 subjects met criteria (Supplemental Figure 1). Replicate injections for unique specimens were averaged, and paired specimens from different areas of the ascending aorta were compared (Figure 4A). Peak and plateau pressures showed that the greater curvature was more vulnerable to initiation and propagation of delamination than the lesser curvature (Figure 4B). There was no difference between proximal and distal zones (Figure 4C), nor between the belly (most dilated) and neck (least dilated) of aneurysmal/ectatic aortas (Figure 4D). By contrast, pressures required for extravasation did not differ among aortic regions, though fewer injections at low flow rate and a lesser incidence of extravasation yielded a low number of events (< 6) in some subgroups, thus limiting meaningful comparisons (Supplemental Figure 2A–C). Variation in peak and plateau pressures among multiple injections in the same region, specimen, or subject was less than half the variation in all regions, specimens, or subjects, supporting the reproducibility of our methods (Supplemental Figure 2D and E). These data confirm that the ex vivo hydraulic fracture model can be used to quantitatively assess vulnerability of the aorta to structural failure and demonstrate that consistent differences exist along circumferential but not axial directions of the ascending aorta.

**Figure 4:**
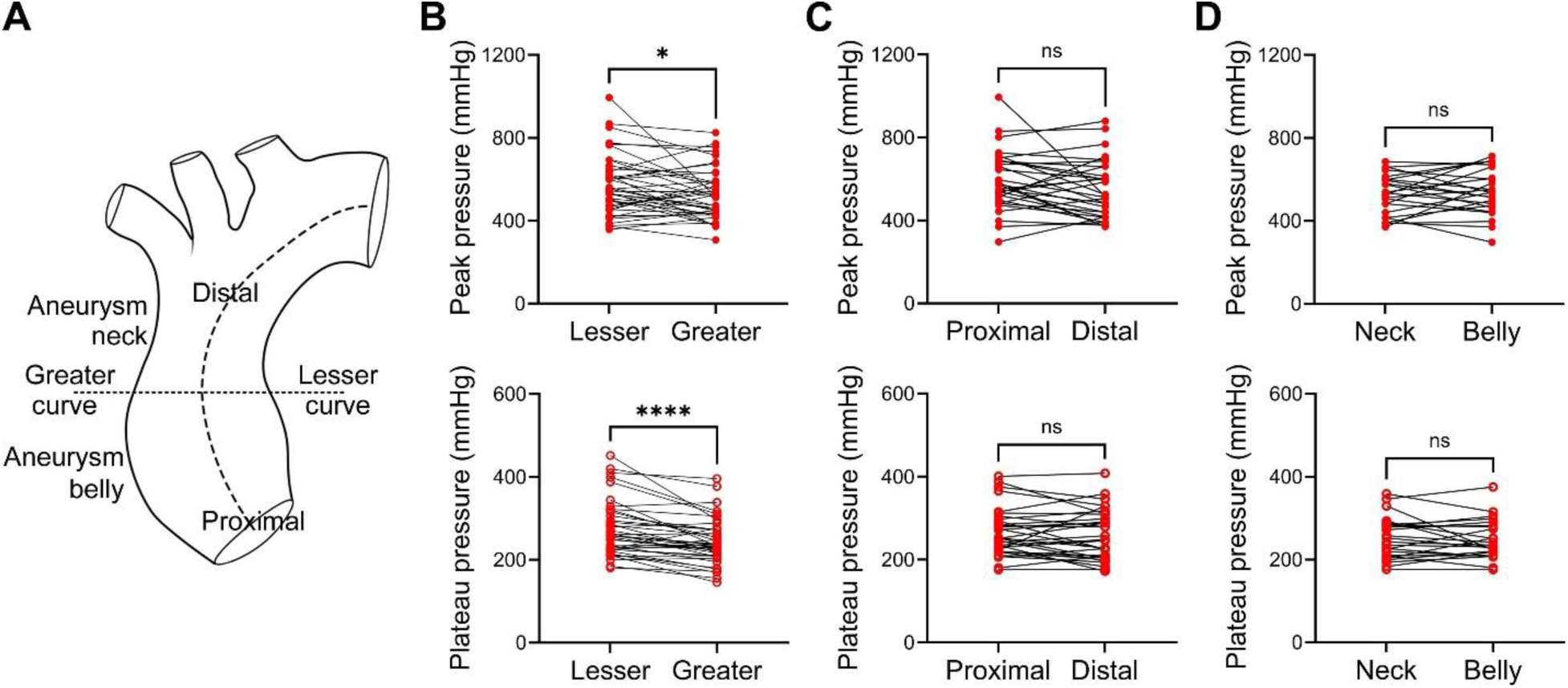
Regional heterogeneity in vulnerability to medial delamination. Peak and plateau pressures required to initiate and propagate medial delamination at high flow rate were compared across different regions of the ascending aorta using paired specimens from subsets of subjects. (**A**) Regional comparisons were proximal vs. distal, lesser vs. greater curvature, and, in dilated aortas, belly (most dilated) vs. neck (least dilated). (**B**) Peak and plateau pressures at the greater curvature were significantly less than at the lesser curvature (*n* = 41). There were no significant differences in pressures between (**C**) proximal and distal specimens (*n* = 31) or (**D**) neck and belly specimens (*n* = 24). Data are shown with lines connecting paired specimens from unique subjects; ns: not significant, \**p* < 0.05, \*\*\*\**p* < 0.0001, Wilcoxon matched-pairs signed rank test (panel B), and paired t-test (panels C and D).

### Age and aortic diameter correlate with vulnerability to medial delamination

We utilized delamination pressures to quantify the impact of clinical risk factors for aortic dissection with numerical variables, namely age and aortic diameter. Peak and plateau pressures required to initiate and propagate medial delamination at high flow rate injection were compared among 90 specimens of 56 subjects (Supplemental Figure 1). Aging correlated with vulnerability to medial delamination, exhibiting significantly lower peak and plateau pressures beyond 70 years of age (Figure 5A and B). Aortic dilatation also correlated with vulnerability to medial delamination, with lower peak and plateau pressures in aortic specimens ≥ 4 cm in diameter (Figure 5C and D). Similar relationships were found when the data were analyzed by individual subjects instead of specimens (Supplemental Figure 3A–D). Normalized measures of aortic diameter have also been used as surgical criteria; Z-score normalizes subject aortic diameter to age-, sex-, and body surface area-matched cohorts (20), while aortic height index is the ratio of subject aortic area to height (21). As with diameter alone, both Z-score and aortic height index correlated with vulnerability to medial delamination but with similar predictive value (Supplemental Figure 4A and B). Interaction analysis by two-way ANOVA showed no significant interaction between the independent variables of age and aortic diameter for the dependent variables of peak or plateau pressures for delamination (Supplemental Figure 4C and D). Extravasation pressures did not vary by age or aortic diameter (Supplemental Figure 5A–D). Despite significant associations of age and aortic diameter with vulnerability to medial delamination, there was substantial overlap of outcomes—many young or non-dilated aortas delaminated at lower pressures than did older or dilated aortas. These results validate age and aortic dilatation as important susceptibility markers for aortic dissection, with additive not interactive effects, but also underscore the influence of factors other than age and diameter.

**Figure 5:**
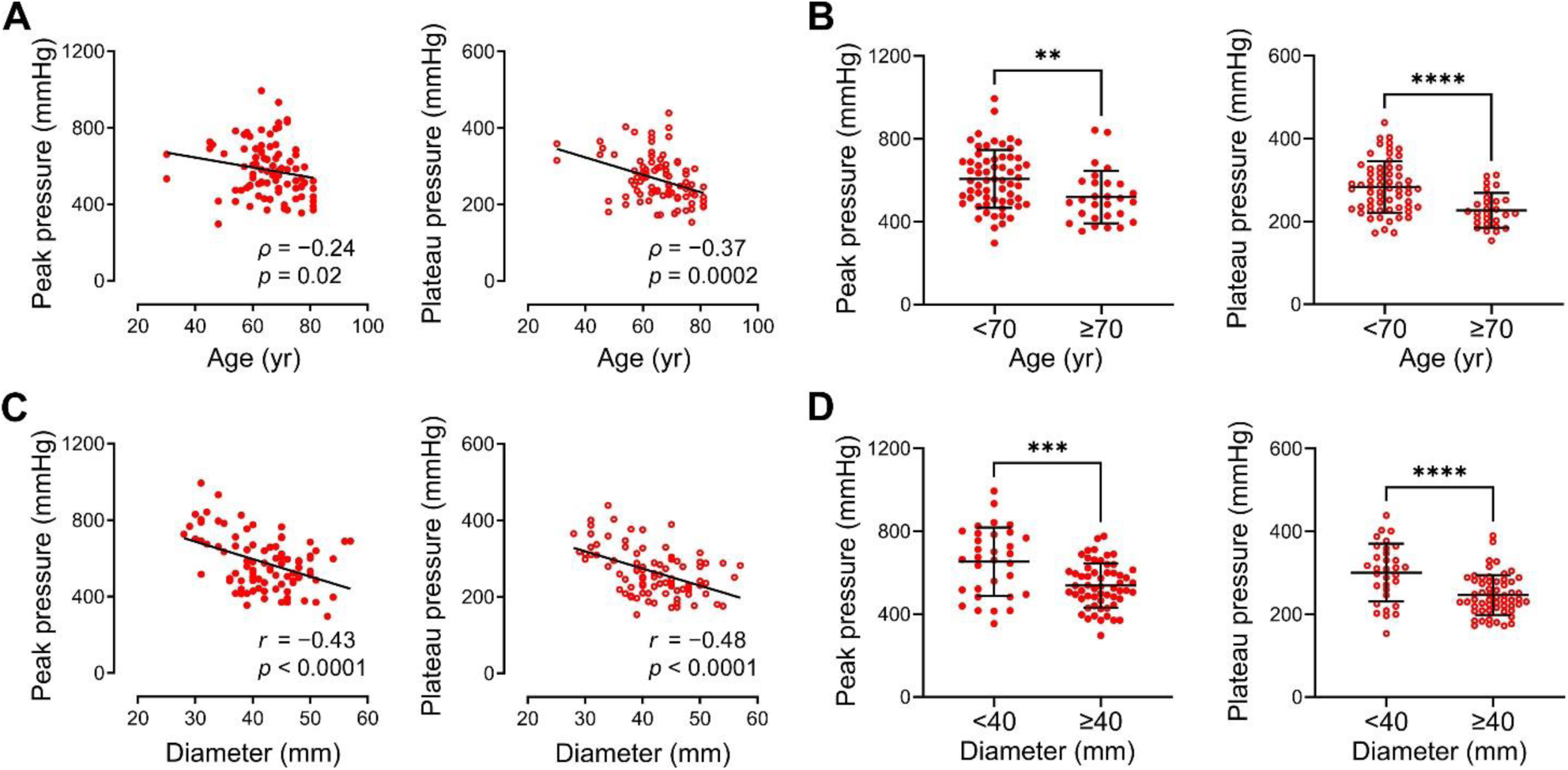
Vulnerability to medial delamination increases with age and aortic dilatation. Peak and plateau pressures required to initiate and propagate medial delamination at high flow rate in ascending aortas from organ donors and surgical patients were compared to age and maximal ascending aortic diameter. (**A**) Increasing age correlated with lower peak and plateau pressures, with (**B**) aortas from subjects ≥ 70 years old having significantly lower peak and plateau pressures. (**C**) Increasing aortic diameter correlated with lower peak and plateau pressures, with (**D**) aortas with diameter ≥ 40 mm having significantly lower peak and plateau pressures. Data for 90 individual specimens are shown (*n* = 62 for age < 70 yr + *n* = 28 for age ≥ 70 yr, *n* = 32 for diameter < 40 mm + *n* = 58 for diameter ≥ 40 mm), \*\**p* < 0.01, \*\*\**p* < 0.001, \*\*\*\**p* < 0.0001, *ρ* correlation coefficient and *p*-value by Spearman correlation with lines indicating best-fit linear regression (panel A), Mann-Whitney test (panel B), *r* correlation coefficient and *p-*value by Pearson correlation with lines indicating best-fit linear regression (panel C), and unpaired t-test (panel D).

### Family history of thoracic aortic disease increases vulnerability while diagnosis of hypertension increases resilience to medial delamination

Expanding on our goal of quantifying subject-specific factors that influence loss of aortic structural integrity, we examined additional demographic and clinical details with categorical variables for associations with delamination pressures in 90 specimens of 56 subjects (Supplemental Figure 1). There was no difference in peak pressure by sex or race and ethnicity (Figure 6A and B). By contrast, family history of thoracic aortic aneurysm or dissection associated with lower peak pressure indicating greater vulnerability for medial delamination (Figure 6C). Interestingly, a diagnosis of hypertension, a well-established risk factor for thoracic aortic aneurysm and dissection, associated with higher peak pressure indicating greater resilience to medial delamination (Figure 6D). Hyperlipidemia, diabetes mellitus, smoking, and number of aortic valve leaflets did not impact peak delamination pressures (Figure 6E–H). When analyzed by individual subjects instead of multiple specimens, the differences in peak pressures for family history and hypertension were no longer significant, though of similar magnitude (Supplemental Figure 6A–H). Plateau pressures did not differ for any of these variables (Supplemental Figure 7A–H). To summarize, familial history and hypertension have opposing influences on susceptibility to medial delamination, although the differences are relatively small.

**Figure 6:**
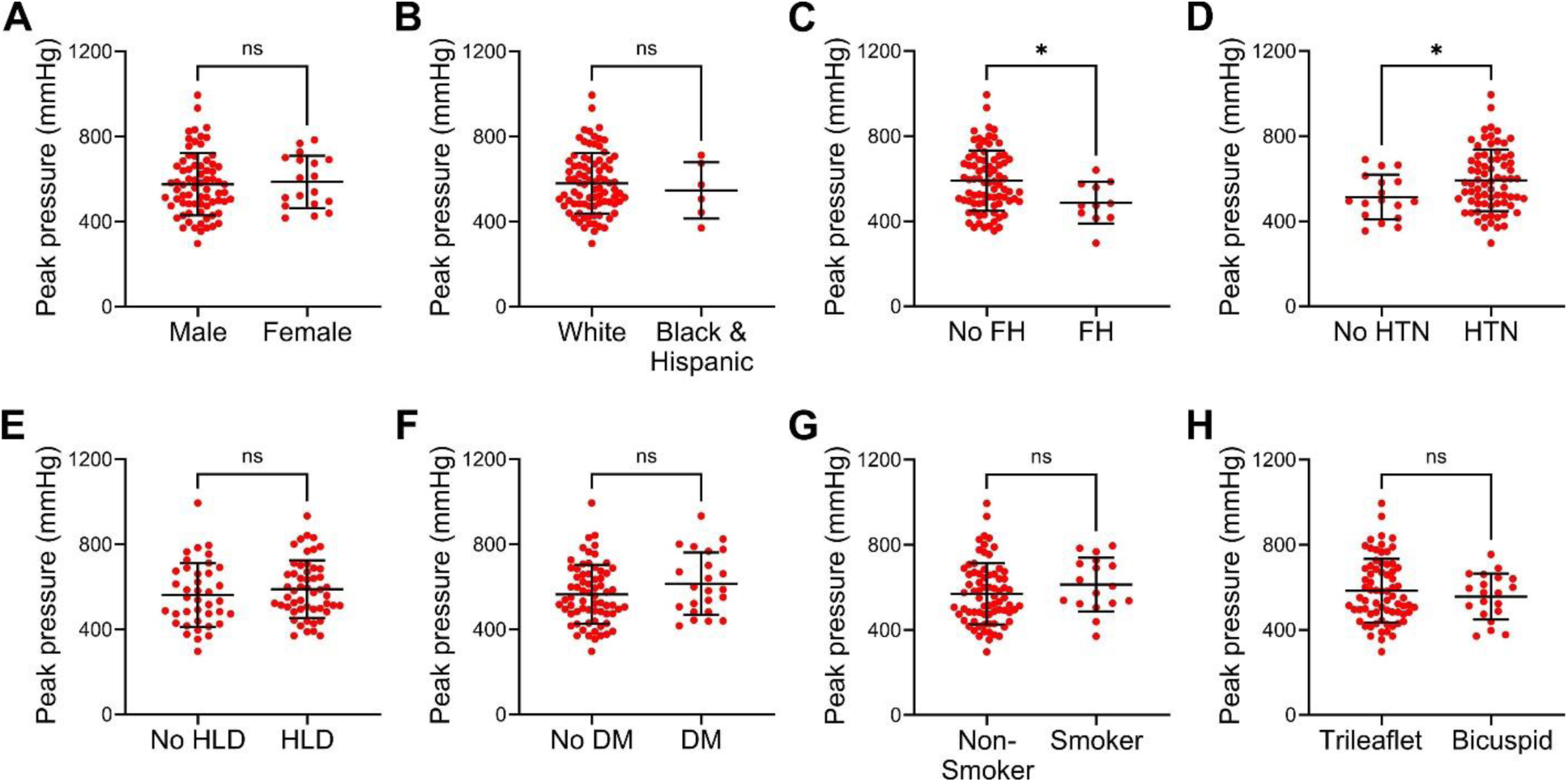
Demographic and clinical determinants of peak pressures required for medial delamination. Peak pressures required to initiate medial delamination at high flow rate in ascending aortas from organ donors and surgical patients were compared by demographic and relevant clinical variables. There was no difference in peak pressures among (**A**) male vs. female subjects or (**B**) white vs. black and Hispanic subjects. (**C**) Subjects with a family history (FH) of thoracic aortic aneurysm or dissection had lower peak pressures, while (**D**) subjects with a diagnosis of hypertension (HTN) had higher peak pressures. (**E**) Hyperlipidemia (HLD), (**F**) diabetes mellitus (DM), (**G**) smoker status, and (**H**) bicuspid aortic valve had no impact on peak pressures. Data for 90 individual specimens are shown (*n* = 72 male + *n* = 18 female, *n* = 84 white + *n* = 6 black and Hispanic, *n* = 80 no FH + *n* = 10 FH, *n* = 17 no HTN + *n* = 73 HTN, *n* = 38 no HLD + *n* = 52 HLD, *n* = 67 no DM + *n* = 23 DM, *n* = 73 non-smoker + *n* = 17 smoker, and *n* = 71 trileaflet + *n* = 19 bicuspid), ns: not significant, \**p* < 0.05, unpaired t-test.

### Greater cell density correlates with vulnerability, while greater collagen and glycosaminoglycan fractions correlate with resilience to medial delamination

To obtain insight into local microstructural factors that may predispose to aortic dissection, we quantified cellular and extracellular matrix components in the undamaged media adjacent to the major delamination plane and examined for associations with delamination pressures. The density of medial cells and fractions of cytoplasm, collagen, elastin, and glycosaminoglycans were determined from histological stains in longitudinal sections of 201 individual injection lesions with delamination at high flow rate (Supplemental Figure 1). Strikingly, increased collagen fraction associated with higher peak and plateau pressures, implying protection from medial delamination (Figure 7A). Conversely, increased cell density associated with lower peak and plateau pressures, indicating vulnerability to medial delamination (Figure 7B). Similar to cell density, cytoplasm fraction calculated from a different histological stain inversely correlated with peak and plateau pressures (Supplemental Figure 8A). Elastin fraction showed no correlation with delamination pressures, whereas greater glycosaminoglycan fraction weakly associated with higher peak but not plateau pressures (Supplemental Figure 8B and C). With regards to physical dimensions, medial thickness showed a positive correlation with plateau pressures, while injection depth, measured as the ratio of the distance from intima to major delamination plane versus total medial thickness, showed a negative correlation with plateau pressures, indicating more effective propagation of delaminations in thinner media or with deeper injections (Supplemental Figure 8D and E). In summary, correlations of local medial elements to delamination pressures show that certain histological hallmarks of medial degeneration, namely loss of smooth muscle cells, medial fibrosis, and mucoid accumulation associated with resilience to medial delamination (Figure 7C). Although loss of elastin did not correlate with delamination pressures, more subtle abnormalities of elastin breaks or disorganization were not assessed. Moreover, one feature of medial degeneration may be a surrogate for another.

**Figure 7:**
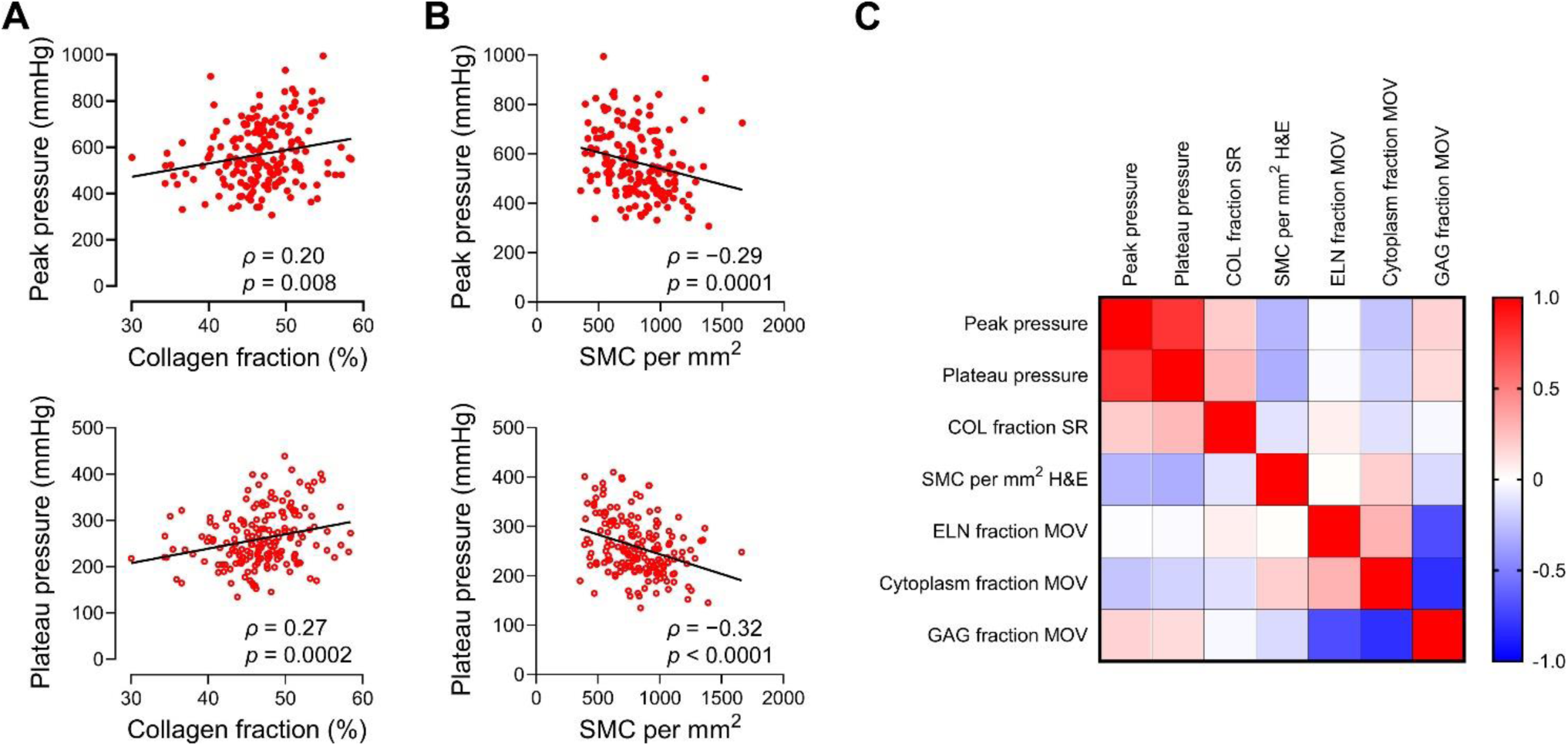
Histological determinants of peak and plateau pressures required for medial delamination. Quantitative histological measurements of medial components in undamaged aorta, immediately adjacent to injection lesions, correlated with the peak and plateau values of pressure required to initiate and propagate medial delamination at high flow rate. (**A**) Collagen fraction associated with higher peak and plateau pressures. (**B**) Smooth muscle cell density showed an inverse relationship with peak and plateau pressures. (**C**) Heatmap matrix of correlation coefficients for delamination pressures and local histological features. Data for individual injection lesions (*n* = 173–183) are shown, depending on suitable quality of histological stains for image quantification. Shown too are *ρ* correlation coefficient and *p*-value by Spearman rank correlation with lines indicating best-fit linear regression (panels A and B), correlation matrix for Spearman rank correlation coefficients (panel C). COL: collagen, SMC: smooth muscle cells, ELN: elastin, GAG: glycosaminoglycan, SR: sirius red, H&E: hematoxylin and eosin, MOV: Movat.

### Decreased cell density in aged aortas and decreased collagen fraction in dilated aortas

To examine if clinical factors and/or histological features are interrelated, we assessed relationships among selected variables that associated with vulnerability or resilience to medial delamination. Histological features of global tissue remodeling were averaged from multiple microscopic fields (3 inner, 3 middle, and 3 outer media) in transverse sections of 90 non-injected specimens serving as references for adjacent injected specimens that delaminated at high flow rate in 56 subjects (Supplemental Figure 1). Age did not correlate with aortic diameter, despite a well-established positive association in populations without aortic disease, likely because of the inclusion of surgical patients with dilated aortas (Figure 8A). Smooth muscle cell density and collagen fraction were independent of each other (Figure 8B). Aging showed a negative correlation with smooth muscle cell density but no association with collagen fraction, whereas aortic dilatation was not associated with smooth muscle cell density but showed a negative correlation with collagen fraction (Figure 8C and D). Thus, aortic remodeling related to aging and aortic dilatation support a resilience role for increased collagen fraction but not decreased cell density. On the other hand, a diagnosis of hypertension associated with increased collagen fraction and decreased smooth muscle cell density supporting a resilience role for both (Figure 8E). When relative amounts of medial components were converted to total amounts per cross-section based on medial thickness and aortic diameter (assuming uniform properties of a hollow cylinder), there were positive correlations between medial thickness and medial cross-sectional area and between the number of smooth muscle cells and collagen content (Supplemental Figure 9A and B). Aging and hypertension correlated with decreased number of smooth muscle cells, but aortic dilatation correlated with increased number of smooth muscle cells and collagen content indicating net gains with aneurysmal remodeling (Supplemental Figure 9C–E). These data indicate complex interconnections among clinical and histological variables and a need to consider both relative and absolute amounts of medial components.

**Figure 8:**
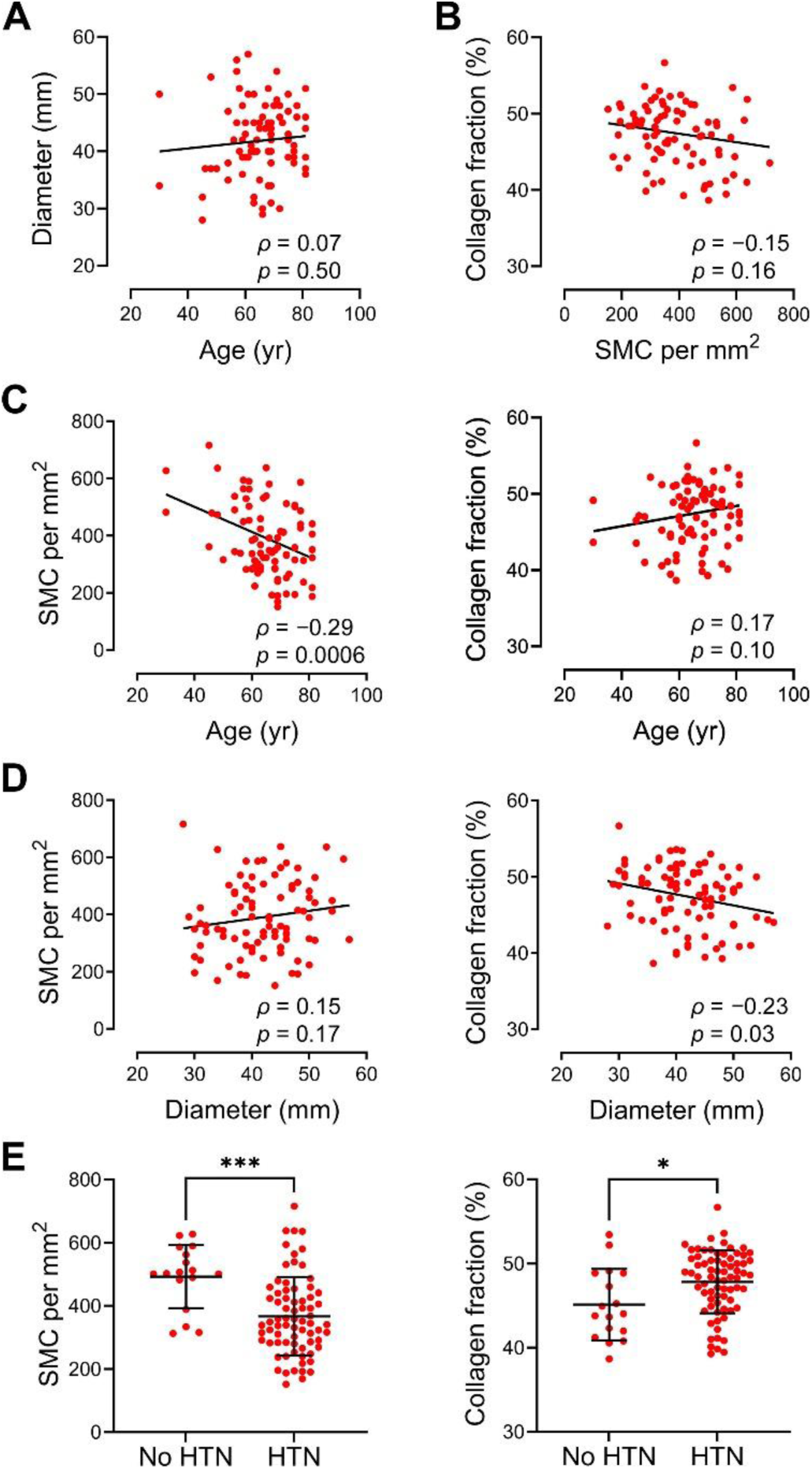
Relationship of clinical factors and relative amounts of medial components. Medial cell density and collagen fraction were averaged from multiple microscopic fields of non-injected ascending aortic specimens serving as controls for adjoining injected specimens that delaminated at high flow rate. (**A**) Aortic diameter was independent of age. (**B**) Collagen fraction was independent of smooth muscle cell density. (**C**) Smooth muscle cell (SMC) density but not collagen fraction was inversely related to age. (**D**) Collagen fraction but not smooth muscle cell density was inversely related to aortic diameter. (**E**) Smooth muscle cell density and collagen fraction were decreased with a diagnosis of hypertension. Data for 90 individual specimens are shown along with *ρ* correlation coefficient and *p*-value by Spearman correlation with lines indicating best-fit linear regression (panels A–D), and unpaired t-test (panel E).

### Increased collagen cross-linking or content, but not acute loss of smooth muscle cells, enhance resilience to medial delamination

To determine if medial cells or fibrillar matrix affect the delamination of elastic lamellae, we pretreated specimens of non-dilated aortas with glutaraldehyde (for protein cross-linking), collagenase (for enzymatic digestion of collagen), or SDS (for detergent lysis of cells) during tissue culture and then compared injection pressures at high flow rate to that of paired untreated controls. Cultured aortic specimens were obtained from 18 organ donors, including 7 subjects that also provided fresh aortic specimens and 11 unique subjects (Supplemental Table 2 and Supplemental Figure 1). Due to varying outcomes of delamination and extravasation with different pressure tracings, we defined failure pressure as the highest value induced during fluid injection. Specimens pretreated with 0.1% glutaraldehyde for 20 hours demonstrated higher failure pressure (exceeding 1000 mmHg in some cases) resulting in either delamination or extravasation (Figure 9A). Histology exhibited no overt changes of the media with and without pre-treatment with fixative (Supplemental Figure 10A and B). By contrast, pretreatment with collagenase A at 1.5 mg/mL for 12 to 24 hours resulted in lower failure pressure with medial injury also manifesting as delamination or extravasation (Figure 9B). Histological analysis of collagenase-treated specimens showed substantial medial erosion with heterogeneous loss of collagen most pronounced in remaining superficial laminae (Supplemental Figure 10C and D). Cell lysis by incubation in 0.5% SDS for 24 to 48 hours did not alter failure pressures or injury pattern as delamination occurred consistently (Figures 9C). Detergent-treated specimens showed loss of nuclear and cytoplasmic features but not extracellular matrix (Supplemental Figure 10E and F). These data confirm that greater extracellular matrix cross-linking and increased collagen content protect against medial delamination but do not reveal a direct role for smooth muscle cell loss.

**Figure 9:**
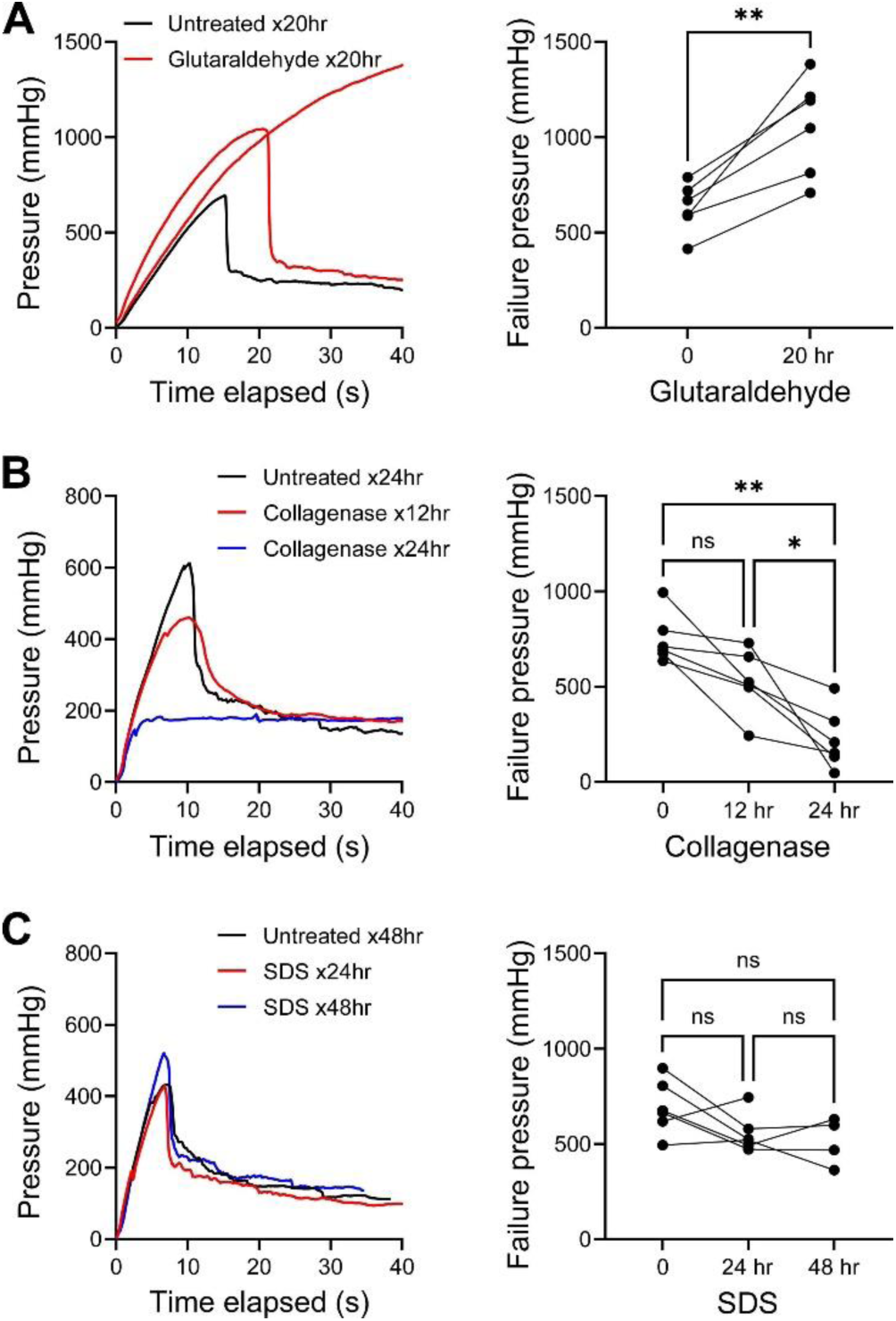
Greater collagen cross-linking or content enhances resilience to medial delamination. Ascending aortic specimens pretreated with various agents or adjacent tissue pieces that remained untreated as controls were injected with fluid at high flow rate and the failure pressure for delamination or extravasation was defined as the highest pressure during injection. (**A**) Representative pressure histories in paired specimens from a single subject and failure pressures in specimens of multiple subjects following protein cross-linking by 0.1% glutaraldehyde at 4 °C for 20 hr vs. untreated controls for 20 hr. (**B**) Representative pressure histories in paired specimens from a single subject and failure pressures in specimens of multiple subjects following collagen digestion by 1.5 mg/mL collagenase A at 37 °C for 12 or 24 hr vs. untreated controls for 24 hr. (**C**) Representative pressure histories in paired specimens from a single subject and failure pressures in specimens of multiple subjects following acute cell lysis by 0.5% SDS at 37 °C for 24 or 48 hr vs. untreated controls for 48 hr. Data for individual specimens are shown with lines connecting paired specimens from unique subjects; (*n* = 6 each, except that specimens from 2 subjects were treated with SDS for 24 but not 48 hr), ns: not significant, \**p* < 0.05, \*\**p* < 0.01, paired t-test (panel A), one-way ANOVA with repeated measures and Tukey’s multiple comparisons test (panel B), mixed-effects analysis with Tukey’s multiple comparisons test (panel C).

## Discussion

In this study, we adapted a previously described method of intramural fluid injection to quantitatively compare subject and structural factors related to vulnerability of the ascending aorta to dissection. We find both a novel mode of medial injury resulting from hydraulic fracturing, termed extravasation, and that some previously described risk factors for aortic dissection, namely a diagnosis of hypertension and certain histological features of medial degeneration, instead correlate with resilience to delamination. These associations are limited by interrelated variables common to aortic remodeling. Mechanistic experiments verify that greater collagen strength and content reduce medial vulnerability to structural wall failure.

Forcing fluid into the aortic wall results in two primary modes of medial damage distinguished by gross and histological appearances as well as by injection pressure tracings. Extravasation is characterized by diffuse minor separation of elastic lamellae, interspersed among areas without separation, with corresponding disruption of medial cells and fibrillar extracellular matrix. By contrast, delamination is characterized by marked separation of elastic lamellae along a single plane with damage limited to a few adjacent laminae. We avoid describing mild-to-moderate separation of elastic lamellae in extravasation lesions as minor delamination to avoid confusion with gross splitting of the media recognized as major delamination. While ex vivo medial delamination is similar to the false lumen well-described in clinical specimens, extravasation resembles ill-defined hemorrhagic lesions evident within the media adjacent to dissection channels (22). The requirement herein for exceedingly elevated pressures to provoke extravasation or delamination supports our recent characterization of healed intimomedial tears in clinical specimens without evidence of medial dissection (23). The spectrum of lesions suggests a multiphasic model of aortic wall failure with sequential initiation and propagation of aortic tears. Moreover, vulnerability to different types of medial damage can be quantified by pressure measurements during intramural fluid injection.

The occurrence of medial delamination versus extravasation differs based on injection flow rate, subject age, and aortic diameter. Prior studies have suggested that elevated concentrations of mechanical stress may contribute to the development of aortic dissection (24, 25). In our fluid injection model, the inserted needle crosses approximately 15 to 20 laminae, based on an inner diameter of 0.3 mm and an interlamellar distance of ∼15 µm (26), hence focusing attention on the substance of the medial wall rather than the intimomedial space where entry tears are expected. Associated splitting of the media between two adjacent elastic lamellae exemplifies the principle of stress concentration as opposed to medial extravasation where many laminae display similar extents of separation and damage. Based on a simple pressure-flow relation (Δ𝑃 = 𝑄𝑅, where pressure drop Δ𝑃 drives flowrate 𝑄 against resistance 𝑅), one would expect a higher pressure if flow is higher and resistance is similar. Hence, higher injection rates can raise local pressure rapidly as seen in the characteristic pressure history and thereby intensify the stress concentration. By contrast, lower injection rates may mitigate stress concentrations, resulting in increased propensity to extravasation. Increasing age and aortic diameter are known clinical risk factors for aortic dissection but also associate with increased aortic stiffness (27, 28). Though more precise work is needed, local heterogeneities in stiffness that accumulate with aging or aortic dilatation may increase stress concentrations, thereby increasing the propensity toward delamination provided that the wall is vulnerable due to normal or sub-normal strength.

We also sought to quantify regional heterogeneity – proximal versus distal, greater versus lesser curvature – in vulnerability of the ascending aorta to medial delamination. Regional differences may result from innate differences in embryological origin of smooth muscle cells or from differences in luminal flow, pressure-dependent distension, or cardiac-dependent extension of the aorta (29–31). Clinical studies suggest that entry tears occur more commonly in the proximal than distal ascending aorta and more commonly in the greater than lesser curvature (32, 33). A prior study using uniaxial peeling found lower delamination strength of the greater than lesser curvature (34). We also show increased vulnerability to medial delamination of the greater versus lesser curvature of the ascending aorta, though with values more consistent for plateau pressures (reflecting propagation) than peak pressures (reflecting initiation). While others have described the ascending aorta as more resilient to delamination than arch or descending thoracic segments (15), we find no difference in vulnerability of the proximal versus distal ascending aorta or of the belly versus neck of aneurysmal/ectatic segments. By performing intramural fluid injections in unpressurized specimens without hemodynamic loads, structural differences were isolated from wall stresses that may modify medial delamination. We conclude that differences in vulnerability of the greater versus lesser curvature are due to differences in medial structure or selective accumulation of medial defects. Conversely, an increased propensity for dissection of the proximal aorta in vivo may reflect the influence of hemodynamics or biaxial wall stresses.

Quantitative testing by fluid injection demonstrates that increasing age and aortic diameter associate strongly with increased vulnerability to delamination. Though these associations are known from prior fluid injection studies in non-dilated aortas or epidemiological studies in subjects with dilated aortas (4, 5, 15, 16, 35), our comparison of dilated and non-dilated aortas indicates independent, additive effects of age and diameter. The increase in vulnerability with age is not negligible; comparisons suggest that a 10-year increase in age results in an average decrease of peak pressures equivalent to a 3 mm increase in diameter and an average decrease in plateau pressures equivalent to a 5 mm increase in diameter. Yet, significant variability in the association of age or aortic diameter and delamination pressures reiterates the influence of additional factors on aortic vulnerability. Other factors including male sex, hypertension, smoking, and family history of thoracic aortic disease have been shown to associate with increased frequency of aortic dissection in epidemiological studies (4, 5, 35). We find no difference in resilience against medial delamination by sex or smoking status, though the study was not intended to be powered for such comparisons. Positive family history of thoracic aortic aneurysm or dissection associated with increased vulnerability, as expected. Surprisingly, hypertension associated with mildly increased resilience (namely, higher peak pressures) to delamination in our study, contrary to population-based studies of risk factors (4, 5, 35). As hypertension is associated with aortic wall remodeling (36), this result suggests that chronic structural changes induced by hypertension may offer mild protection from aortic dissection while acutely increased hemodynamic stresses from blood pressure elevation (e.g., vigorous exercise) can trigger dissection events. Additional tests of loaded, pressurized aortic specimens would better determine relationships between structural properties and hemodynamically induced stresses.

Medial degeneration, characterized by elastin loss and fragmentation, accumulation of mucoid material (glycosaminoglycans), fibrosis (collagen deposition), and smooth muscle cell loss, has long been associated with thoracic aortic aneurysms (37–39). Other groups, and more recently our group, have emphasized that degenerative changes are also related to aging and that such changes need not imply vulnerability to dissection (26, 40, 41). Notably, in experimental studies of Hutchinson-Gilford progeria syndrome, the aorta has tremendous loss of smooth muscle cells and enormous diffuse accumulation of proteoglycans yet does not dissect (42). By comparing injection pressures and local histological parameters adjacent to delamination lesions, we found that certain hallmarks of medial degeneration—smooth muscle cell loss and increased collagen—associated strongly with increased resilience to delamination while another, diffuse accumulation of proteoglycans, was associated with resilience mildly. Elastin fraction, however, had no significant effect on pressures required for delamination. These observations suggest that some features of the broad descriptor of medial degeneration, particularly diffuse accumulations of collagen and perhaps glycosaminoglycans, may represent reparative responses of smooth muscle cells in areas of medial damage, with partially adaptive effects conferring greater resilience to delamination. Yet, local deposits instead of diffuse accumulation may lead to stress concentration and greater aortic vulnerability (43).

Further measurement of global histological changes in non-injected control specimens revealed interrelated variables with aortic remodeling. Increasing age correlated with lower smooth muscle cell density, a contradictory association given lower delamination pressures in older subjects but higher delamination pressures in specimens with lower cell density. A diagnosis of hypertension associated with increased collagen fraction and decreased cell density in keeping with resilience roles. In contrast, larger aortic diameter associated with decreased collagen fraction, consistent with the increased vulnerability seen with both aortic dilation and lower collagen fraction. Yet, cross-sectional collagen content and total number of cells increased with aortic dilatation, indicating net gains. We have previously documented medial hypertrophy and cellular hyperplasia in thoracic aortic aneurysms in independent cohorts of 57 and 35 subjects (26, 44). Thus, although the aortic wall thins with aneurysmal enlargement, it thins less than predicted from the degree of aortic dilatation because of increased tissue mass. We suggest that changes in hemodynamic stresses that drive or result from aortic dilatation can induce adaptive responses that increase total collagen content, but when this adaptive response is insufficient to normalize the proportion of collagen within the media it becomes vulnerable to delamination, particularly in local areas with greater loss of collagen.

We performed gain- and loss-of-function experiments to verify if loss of smooth muscle cells or collagen increase vulnerability to medial delamination. Glutaraldehyde, collagenase, and SDS were selected to induce protein cross-linking, collagen digestion, and acute cell lysis, respectively. Aortic specimens pretreated with glutaraldehyde were significantly more resilient to delamination, with some medial injury manifesting as extravasation at extremely high pressures. These findings are consistent with other methods documenting increased resistance to peeling of glutaraldehyde-treated aortas (45). Prolonged incubation of aortic specimens with collagenase resulted in thinning of aortic tissue due to progressive loss of outer laminae, a finding that itself demonstrates the necessity of collagen to hold elastic lamellae together. Thinning of the media prevented homogenous digestion of all medial collagen but we observed a profound decrease in collagen density in the periphery of the remaining tissue. Extravasation as an injury pattern occurred more frequently, and failure after medial injection occurred at significantly decreased pressures, similar to prior studies measuring force required to peel collagenase-treated aortas (45). After SDS-mediated acute cell lysis, we saw no difference in medial injury outcomes or delamination pressures after intramural fluid injection. These data are not consistent with the observation of higher delamination pressures in areas of decreased cellular density, suggesting that interrelated in vivo remodeling events accompanying local loss of smooth muscle cells are responsible for increased resistance to delamination. These mechanistic results support a direct role for collagen resisting separation of elastic lamellae under passive conditions, although a protective role for smooth muscle cell contraction cannot be excluded (46).

In conclusion, medial fluid injection is a reliable method for testing the biomechanical properties of the aortic wall with respect to structural failure. This test is simple, inexpensive, reproducible, and requires little special equipment. Unlike biaxial or uniaxial testing, a single consistent protocol can be used, and there is no need for constitutive modelling before comparison of results (28, 47). Thus, results can easily be compared between different centers. Fluid injections can be conducted in aortas under physiological conditions, including in warmed, oxygenated media with aortas under relevant biaxial stretch, but an advantage of our method on unloaded tissue at room temperature is the isolation of aortic structural properties from induced wall stress or active cellular properties. With respect to clinical risk factors, we identify older age and larger aortic diameter as relatively robust markers and positive family history as a weaker marker for vulnerability, with a diagnosis of hypertension as a comparatively weak marker for resilience to medial delamination. Epidemiological and clinical research should work towards integrating age into surgical guidelines for replacement of the thoracic aorta and unraveling nuanced effects of different levels of chronic hypertension from acute elevations in blood pressure. With respect to medial structure, we note that certain historical features of medial degeneration yet correlate with resistance to delamination. These findings are seemingly contrary to associations between medial degeneration and aging or aortic dilatation and between aging or aortic dilatation and aortic vulnerability. We propose that the accumulation of collagen that frequently occurs in areas of medial damage characterized by elastin breaks, widely interpreted as evidence of structural deterioration, can be reparative responses by smooth muscle cells that reduce vulnerability to aortic dissection. Chronic hypertension resulting in medial fibrosis may increase resilience to aortic dissection by requiring higher pressures for delamination yet still predispose to even greater acute elevations of blood pressure that trigger aortic dissection. A corollary is that a diagnosis of hypertension would be a more significant risk factor for dissection of susceptible aortas if not for this form of hypertensive preconditioning and increased tissue fibrosis.

## Methods

### Study approval

Research protocols to obtain aortic tissue from deceased organ donors were approved by the Yale Institutional Review Board with waiver of consent and by the New England Organ Bank with written consent for research of non-transplanted tissues from the next-of-kin. All procedures were in accordance with federal and institutional guidelines.

### Subjects

Aortic tissue was collected from 76 subjects who underwent elective thoracic aortic surgery or were organ donors whose hearts were not used for transplantation from September 2023 to October 2024. Specimens were procured by the investigators in the operating room to ensure correct anatomical location and orientation. Demographic and clinical information were recorded from review of medical records. Aortic diameters were measured from chest computed tomography scans using multiplanar reconstruction to ensure double-oblique views. Fold-increase and Z-score were determined using a custom calculator that indexes ascending aortic diameter to age, sex, weight, and height (https://medicine.yale.edu/surgery/cardio/research/). Aneurysm was defined as > 1.5-fold expected diameter, ectasia as > 2 SD but ≤ 1.5-fold expected diameter, and non-dilated as ≤ 2 SD of expected diameter. Neck and belly aortic specimens were defined as the most and least dilated zones of ascending aortas with maximal diameter > 4 cm. Aortic valve abnormalities were determined by pre-operative transthoracic or intraoperative transesophageal echocardiography. Family history of thoracic aortic aneurysm or dissection was defined as documented occurrence in first-degree relatives. Genetic testing was performed in 15 of 41 patients undergoing surgery for thoracic aortic aneurysm and 1 pathogenic variant was found in a 26-year-old male of NM_053025.4(MYLK):c.3393dup (p.Thr1132fs).

### Intramural fluid injection of aortic specimens

Resected ascending aortas were divided by the investigators into different regions resulting in one or more specimens per subject. The specimens were stored at 4 °C and tested within 72 hours. Circumferential rings of ascending aorta were oriented and opened along the lesser curvature. Loose perivascular tissue was removed by sharp dissection. Pieces of aorta were secured with pins, intimal side up, and kept submerged in normal saline at room temperature for the entirety of the fluid injection experiment. A 25-gauge butterfly needle (Becton Dickinson) was inserted 1–2 mm into the aortic wall at an angle nearly parallel to the tissue. The needle was connected via high-pressure tubing and a three-way stopcock to a syringe pump and pressure transducer (Edwards Lifesciences, Irvine, CA) connected to a PowerLab device and LabChart software (ADInstruments, Colorado Spring, CO). A syringe pump (Fisher Scientific, Model 780100I) was used to infuse 0.9% saline solution with dilute India ink 1% v/v and 5% human albumin (AlbuRx, CSL Behring). Infusion rates and times were 100 µL/min for 120 s designated as low flow rate or 300 µL/min for 40 s designated as high flow rate, thus resulting in equal volumes of intramural fluid injected.

#### Pretreatment of aortic specimens

After specimen preparation as above, 46 tissue pieces measuring 4 x 2 cm were subjected to various treatments during tissue culture. For protein cross-linking, aortic specimens were incubated with 0.1% glutaraldehyde (Sigma-Aldrich) in phosphate buffered saline (Gibco) at 4 °C for 20 hr (48). For collagen digestion, aortic specimens were incubated with 1.5 mg/mL collagenase A (10103578001, Roche) in DMEM (Gibco) for 12 to 24 hours at 37 °C (12). For cell lysis, aortic specimens were incubated with 0.5% SDS (AmericanBio) in PBS for 24 to 48 hours at 37 °C (49, 50). Control specimens were incubated under the same conditions without fixative, enzyme, or detergent and designated as untreated. Cultured aortic specimens were rinsed with PBS following treatment and tested by intramural fluid injection.

#### Histology and histomorphometry

Transverse sections of aortic specimens were fixed in 10% neutral buffered formalin overnight at 4 °C, transferred to 70% ethanol for 24 hours, and embedded in paraffin blocks. Serial 5 μm-thick sections were stained with hematoxylin and eosin, sirius red, and Movat’s pentachrome stains by Yale’s Research Histology Laboratory using standard techniques and scanned using an Aperio AT2 scanner (Leica). Not all histological stains were of sufficient quality and, initially, not all replicate injection lesions were processed yielding 65 non-injected, control specimens, 63 extravasation lesions at low flow rate, and 201 delamination lesions at high flow rate for analysis. Medial wall thickness and injection depth were measured using QuPath software (https://qupath.github.io) from Movat’s pentachrome stained images. Quantification of local media features was measured in two undamaged areas on either side and adjacent to the major delamination plane of each injection lesion from longitudinally cut slides. Global media features were averaged from 9 fields of the inner, middle, and outer media of non-injected control aortic specimens from transversely cut slides. Smooth muscle cell density and collagen fraction were calculated from hematoxylin and eosin stains and sirius red stains, respectively, using ImageJ software (http://imagej.net). For collagen fraction, the automated thresholding function was used to determine percent area of positive staining. For smooth muscle cell density, images underwent contrast enhancement by splitting into the red channel and thresholding, and nuclei were counted using the Analyze Particles function. Medial fractions (percent area of positive staining) of elastin, cytoplasm, and glycosaminoglycans were measured from Movat pentachrome-stained images using MATLAB software (MathWorks) and a color segmentation algorithm (https://github.com/yale-humphrey-lab/histology-analysis).

#### Immunofluorescence and confocal microscopy

Formalin-fixed, paraffin-embedded aortic specimens were sectioned at 5 μm. The slides were serially deparaffinized in xylene and gradually rehydrated in ethanol and water. After heat-mediated antigen retrieval (H-3300-250, Vector Laboratories), sections were incubated with antibodies to smooth muscle α-actin (clone 1A4-eFluor 570 conjugate, 41-9760-82, Invitrogen) and collagen III (1330-01, Southern Biotech). Secondary labelling of unconjugated primary goat antibody was performed with Alexa Fluor 488-conjugated IgG (Invitrogen). Elastin was labelled with Alexa Fluor 633 hydrazide (A30634, Invitrogen) and DNA with DAPI (D1306, Invitrogen). Images were acquired using a Stellaris 8 Falcon confocal microscope with LAS X software (Leica).

#### Statistics

Categorical data are expressed as counts and percentages, numerical data are expressed as mean ± SD, and individual values are shown in graphs. Data were checked for normality by Shapiro-Wilk test and non-parametric tests were used for variables determined as not normally distributed. Comparison of categorical data was by Fisher’s exact test. Comparison of numerical variables between 2 groups was by Mann-Whitney test or Wilcoxon matched pairs signed rank test for nonparametric data and by unpaired or paired t-test for parametric data. Comparison of numerical variables between multiple groups was by ANOVA or mixed-effects analysis. Strength and direction of associations were determined by Spearman correlation for nonparametric data or by Pearson correlation for parametric data. All tests were performed with Prism 10.4.1 (GraphPad Software). A two-tailed *p*-value < 0.05 was considered to indicate statistical significance.

## Author Contributions

AC, GT, and RA designed the study. JDH and AC designed data acquisition. AC, KW, and DL conducted experiments and acquired data. PV, GT, and RA contributed specimens. AC, GT, and RA analyzed and interpreted data. GT and RA supervised the work. AC, PV, JDH, GT, and RA wrote and edited the manuscript. The two senior authors equally shared supervision of the work.

