## Supplemental Materials for "Increased medial collagen enhances aortic resilience against mural delamination from hydraulic fracturing"

**Supplemental Table 1: Demographics and clinical features of human subjects providing fresh aortic specimens**

| | Organ Donors $n = 24$ | TAA patients $n = 41$ | $p$-value |
|---|---|---|---|
| Age (yr) | 59.9 ± 16.0 | 63.5 ± 11.6 | 0.30 |
| Sex: Female | 8 (33.3%) | 6 (14.6%) | 0.12 |
| Male | 16 (66.7%) | 35 (85.4%) | |
| Race: Black | 3 (12.5%) | 1 (2.44%) | 0.14 |
| White | 21 (87.5%) | 40 (97.6%) | |
| Ethnicity: Hispanic | 1 (4.17%) | 1 (2.44%) | > 0.99 |
| Non-Hispanic | 23 (95.8%) | 40 (97.6%) | |
| Body surface area ($m^2$) | 2.04 ± 0.33 | 2.07 ± 0.18 | 0.75 |
| Ascending aorta diameter (mm) | 36.0 ± 5.1 | 45.8 ± 5.3 | < 0.0001 |
| Ascending aorta fold-increase | 1.05 ± 0.12 | 1.31 ± 0.16 | < 0.0001 |
| Ascending aorta Z-score | 0.50 ± 1.29 | 3.44 ± 1.82 | < 0.0001 |
| Ascending aorta aneurysm | 0 (0%) | 5 (12.2%) | 0.15 |
| Ascending aorta ectasia | 4 (16.7%) | 27 (65.9%) | < 0.0001 |
| Ascending aorta non-dilated | 20 (83.3%) | 9 (22.0%) | < 0.0001 |
| Bicuspid aortic valve | 0 (0%) | 12 (29.3) | 0.002 |
| Aortic valve regurgitation | 0 (0%) | 11 (26.8%) | 0.0048 |
| Aortic valve stenosis | 0 (0%) | 8 (19.5%) | 0.022 |
| Family history of TAAD | 0 (0%) | 7 (17.1%) | 0.041 |
| Hypertension | 19 (79.2%) | 31 (75.6%) | > 0.99 |
| Hyperlipidemia | 13 (54.2%) | 21 (51.2%) | > 0.99 |
| Diabetes mellitus | 9 (37.5%) | 4 (9.76%) | 0.023 |
| Smoker, former or active | 9 (37.5%) | 5 (12.2%) | 0.031 |
| Coronary artery disease | 7 (29.2%) | 6 (14.6%) | 0.12 |

*Study subjects that provided fresh aortic specimens for intramural fluid injection consisted of organ donors without known aortic disease and patients undergoing surgery for thoracic aortic aneurysms. Demographic and clinical data were extracted from the electronic medical records. Maximal diameter of the ascending aorta was measured from chest computed tomography scans and indexed to age, sex, and body size. Some organ donors had ectatic ascending aortas and some patients with root or arch aneurysms had non-dilated ascending segments. Continuous variables are represented as mean ± SD; categorical variables are represented as number of subjects with % subjects in parentheses. Comparisons of continuous variable are by unpaired t-test. Comparisons of categorical variables are by Fisher's exact test. TAAD: thoracic aortic aneurysm and dissection.

**Supplemental Table 2: Demographics and clinical features of human subjects providing aortic specimens for tissue culture**

| | Glutaraldehyde *n* = 6 | Collagenase *n* = 6 | SDS *n* = 6 | *p*-value |
|---|---|---|---|---|
| Age (yr) | 56.8 ± 15.8 | 60.7 ± 8.4 | 67.5 ± 3.0 | 0.23 |
| Sex: Female | 3 (50.0%) | 3 (50.0%) | 2 (33.3%) | > 0.99 |
| Male | 3 (50.0%) | 3 (50.0%) | 4 (66.7%) | |
| Race: Black | 0 (0%) | 1 (16.7%) | 0 (0%) | > 0.99 |
| White | 6 (100%) | 5 (83.3%) | 6 (100%) | |
| Ethnicity: Hispanic | 1 (16.7%) | 0 (0%) | 0 (0%) | > 0.99 |
| Non-Hispanic | 5 (83.3%) | 6 (100%) | 6 (100%) | |
| Body surface area ($m^2$) | 1.89 ± 0.27 | 2.02 ± 0.31 | 2.26 ± 0.34 | 0.15 |
| Aorta diameter (mm) | 34.8 ± 2.9 | 38.0 ± 2.2 | 40.8 ± 6.4 | 0.09 |
| Aorta fold-increase | 1.06 ± 0.07 | 1.10 ± 0.08 | 1.13. ± 0.16 | 0.61 |
| Aorta Z-score | 0.67 ± 0.76 | 1.17 ± 0.93 | 1.63 ± 1.98 | 0.48 |
| Aorta ectasia | 0 (0%) | 0 (0%) | 3 (50.0%) | 0.07 |
| Aorta non-dilated | 6 (100.0%) | 6 (100.0%) | 3 (50.0%) | |
| Hypertension | 4 (66.7%) | 5 (83.3%) | 5 (83.3%) | > 0.99 |
| Hyperlipidemia | 3 (50.0%) | 3 (50.0%) | 2 (33.3%) | > 0.99 |
| Diabetes mellitus | 1 (16.7%) | 1 (16.7%) | 1 (16.7%) | > 0.99 |
| Smoker, former or active | 2 (33.3%) | 4 (66.7%) | 4 (66.7%) | 0.59 |
| Coronary artery disease | 3 (50.0%) | 0 (0%) | 3 (50.0%) | 0.15 |

*Organ donors without known aortic disease provided aortic specimens for tissue culture to pretreat with glutaraldehyde, collagenase, or SDS versus untreated controls. Demographic and clinical data were extracted from the electronic medical records. Maximal diameter of the ascending aorta was measured from chest computed tomography scans and indexed to age, sex, and body size. Some organ donors had ectatic ascending aortas. Continuous variables are represented as mean ± SD; categorical variables are represented as number of subjects with % subjects in parentheses. Comparisons of continuous variable are by one-way ANOVA. Comparisons of categorical variables are by Fisher's exact test.

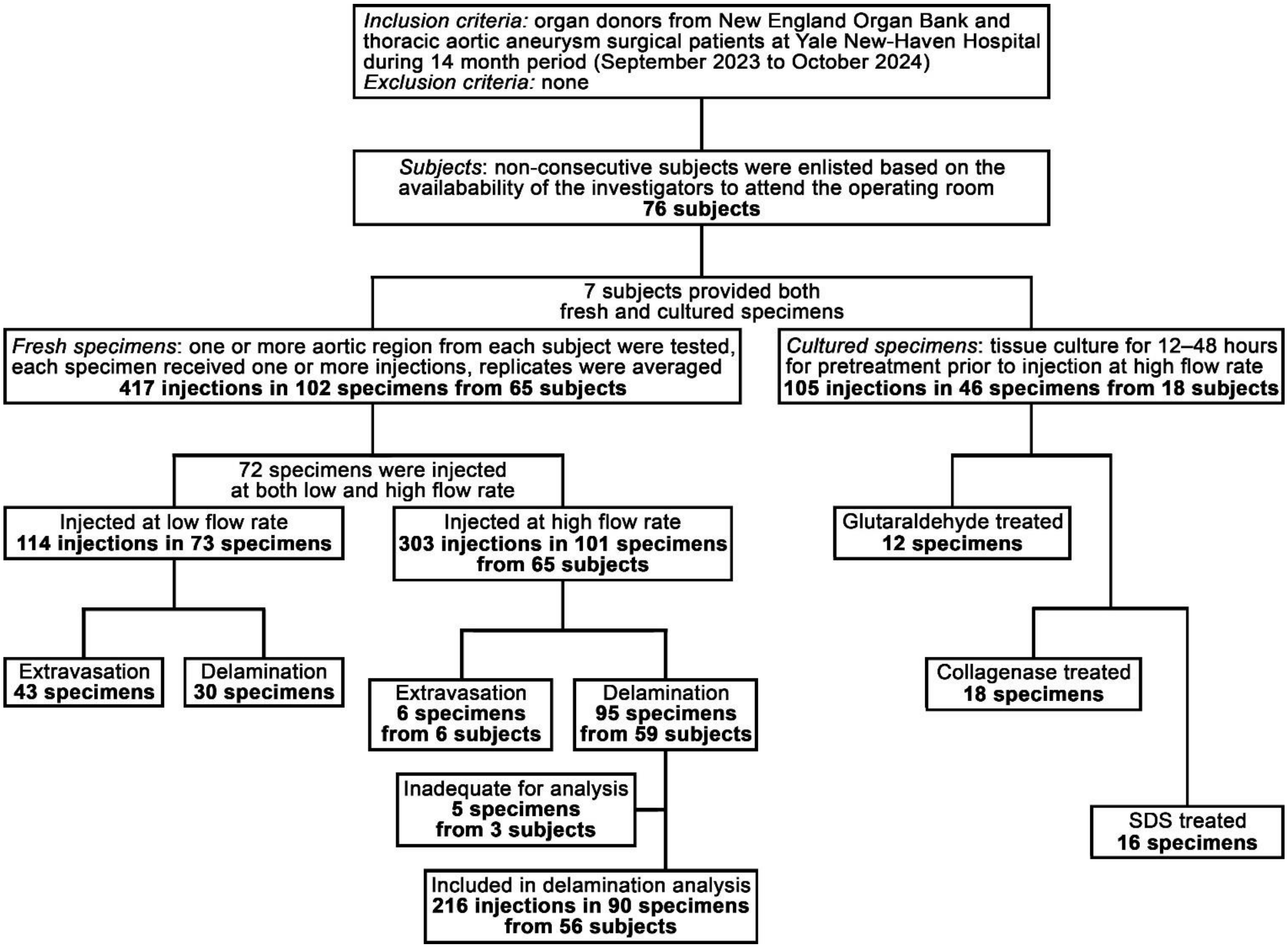

**Supplemental Figure 1. Flow diagram of study design**. Summary of inclusion/exclusion criteria, number of subjects enlisted, and number of specimens injected. The total number of specimens included separate regions of the ascending aorta from some subjects. Single injections, replicate injections of the same region at the same flow rate, or multiple injections in different regions or at different flow rates were performed on individual specimens, ranging from 1–6 injections depending on the size of the specimen, and results from replicate injections were averaged. Including replicate injections, a total of 417 injections in 102 fresh aortic specimens from 65 subjects were completed; 72 specimens were injected at both low and high flow rates via separate injections. Of 303 high flow rate injections in fresh aortic specimens, 28 injections resulted in extravasation, 13 injections resulted in intermediate extravasation/delamination phenotype, and 46 injections with delamination yielded pressure tracings unsuitable for quantification and were excluded from delamination pressure analyses. A final analysis of 90 specimens with delamination at high flow rate were based on 216 successful injections (mean of 2.4 ± 1.3 injections per specimen) derived from 56 of 65 subjects. Besides injection of fresh aortic specimens, 46 specimens from 18 subjects (including 7 subjects that also provided fresh specimens and 11 unique subjects) were cultured for 12–48 hr during glutaraldehyde, collagenase, or SDS treatment prior to fluid injection at high flow rate.

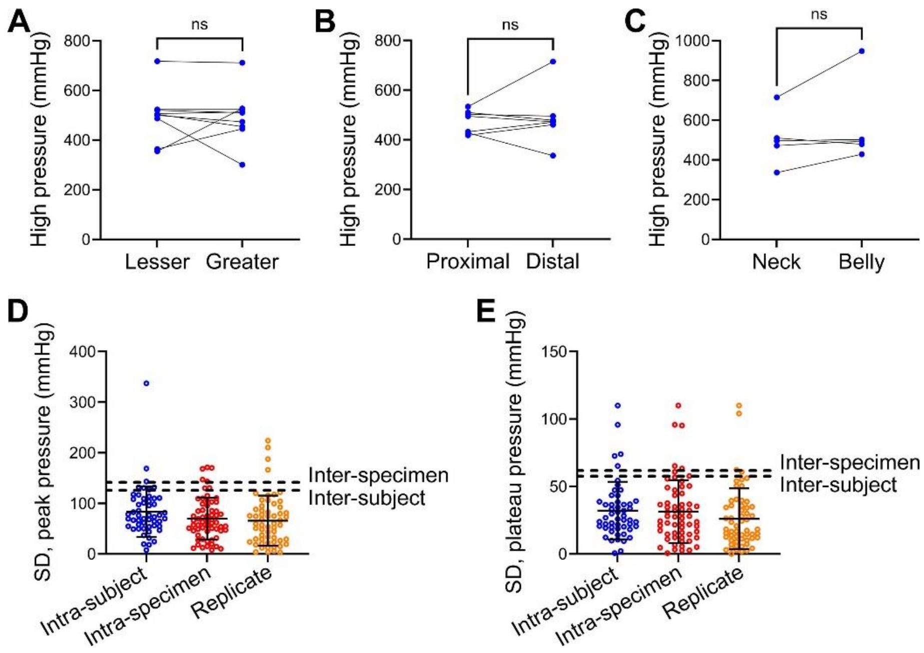

**Supplemental Figure 2. Homogeneity of extravasation pressures in different regions of the ascending aorta and variability in delamination pressures.** Terminal high pressures required for extravasation at low flow rate were compared across regions of the ascending aorta using paired specimens from subsets of subjects. There were no significant differences in extravasation pressures between (**A**) lesser and greater curvature, *n* = 9, (**B**) proximal and distal, *n* = 7, or (**C**) neck and belly specimens, *n* = 5. Data for individual specimens are shown with lines connecting paired specimens from unique subjects; ns: not significant, Wilcoxon matched-pairs signed rank test (panels A–C). Additionally, variation measured as SD of (**D**) peak and (**E**) plateau pressures required for delamination at high flow rate among multiple injections in different specimens of individual subjects (intra-subject variation), among multiple injections in different regions of individual specimens (intra-specimen variation), and among multiple injections in single regions of individual specimens (replicate variation) compared to that among different specimens (inter-specimen variation) and subjects (inter-subject variation). Individual data are shown for intra-subject (*n* = 52), intra-specimen (*n* = 61), and replicate (*n* = 58) injections with lines indicating mean ± SD, while inter-specimen (*n* = 90) and inter-subject (*n* = 56) SD are represented by interrupted lines from data shown in Figure 5 and Supplemental Figure 3.

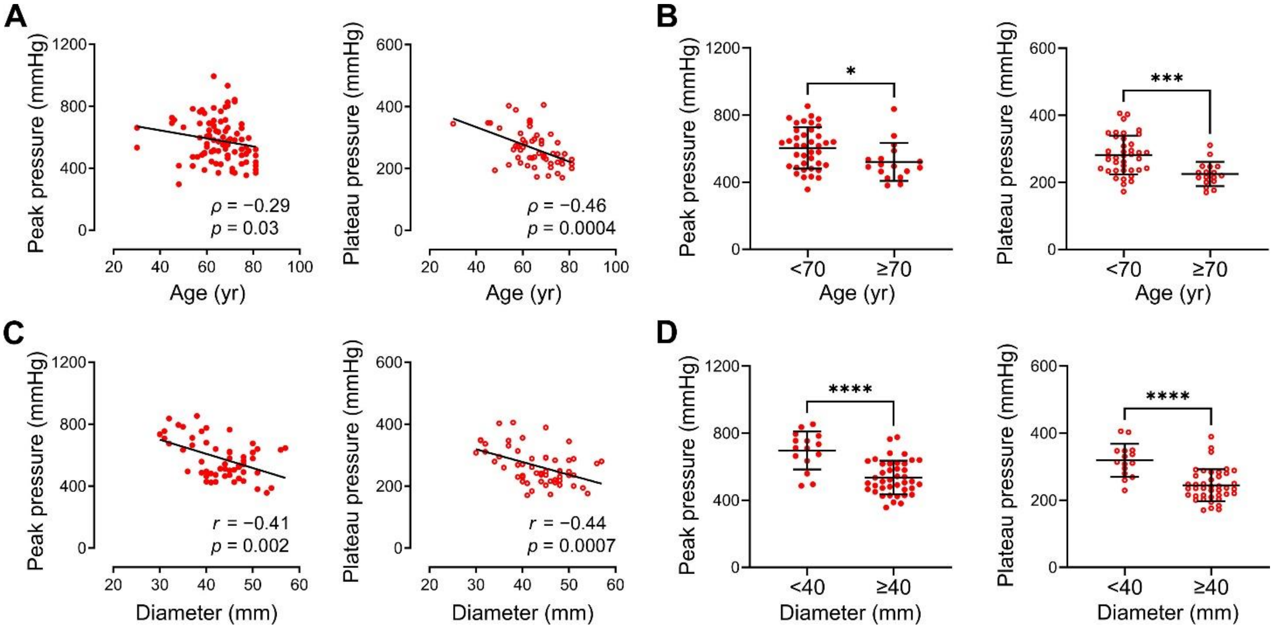

**Supplemental Figure 3: Correlation of age and aortic dilatation to medial delamination pressures by subjects.** Peak and plateau pressures required to initiate and propagate medial delamination at high flow rate in ascending aortas from organ donors and surgical patients were compared to age and maximal ascending aortic diameter. (**A**) Increasing age correlated with lower peak and plateau pressures, with (**B**) aortas from subjects ≥ 70 years old having statistically lower peak and plateau pressures. (**C**) Increasing aortic diameter correlated with lower peak and plateau pressures, with (**D**) aortas of diameter ≥ 40 mm having statistically lower peak and plateau pressures. Data for individual subjects are shown, $n$ = 56 ($n$ = 39 for age < 70 yr + $n$ = 17 for age ≥ 70 yr; $n$ = 15 for diameter < 40 mm + $n$ = 41 for diameter ≥ 40 mm), *$p < 0.05$, ***$p < 0.001$, ****$p < 0.0001$, $\rho$ correlation coefficient and $p$-value by Spearman correlation with lines indicating best-fit linear regression (panel A), Mann-Whitney test (panel B), $r$ correlation coefficient and $p$-value by Pearson correlation with lines indicating best-fit linear regression (panel C), and unpaired t-test (panel D).

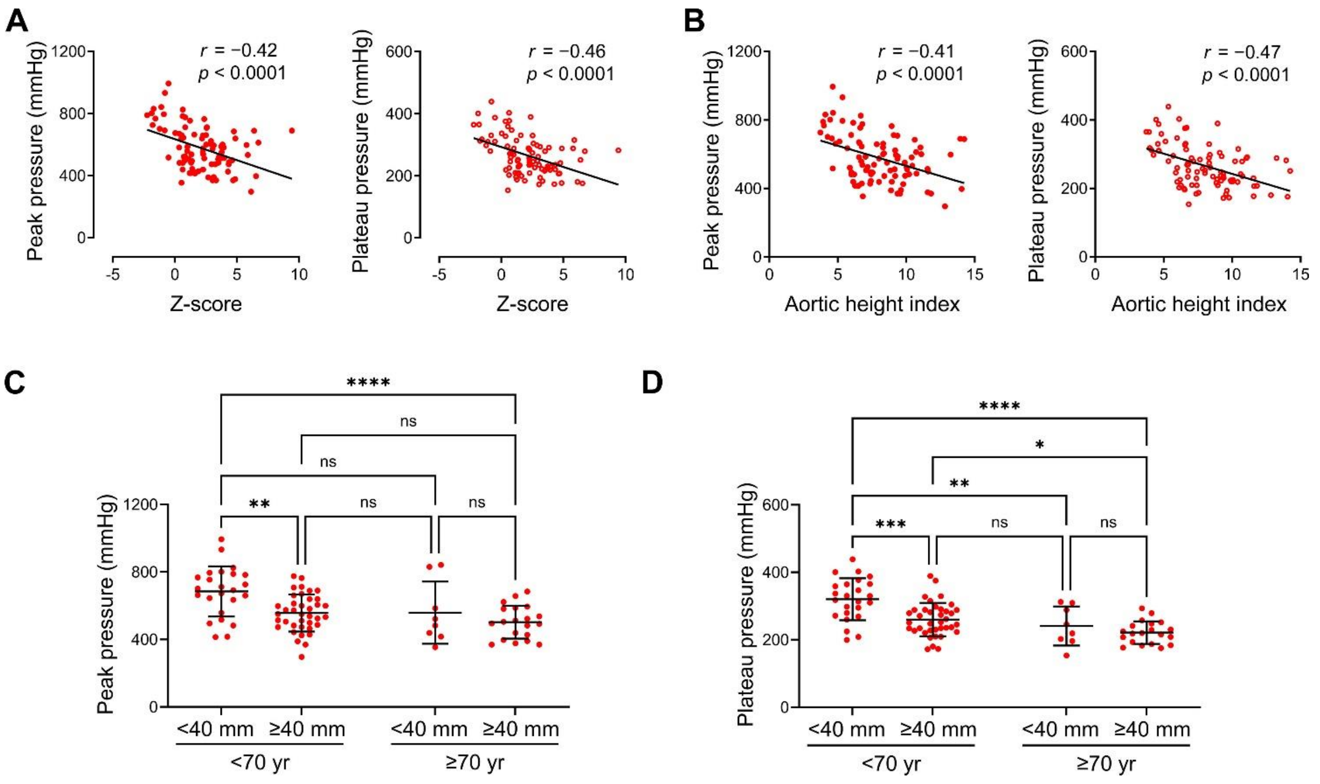

**Supplemental Figure 4: Normalized measures of aortic size and two-way analysis of age and diameter.** (**A**) Z-score and (**B**) aortic height index of maximal ascending aortic diameter demonstrated an inverse relationship with peak and plateau pressures required to initiate and propagate medial delamination, respectively. Data for individual specimens are shown, $n = 90$, $r$ correlation coefficient and $p$-value by Pearson correlation with lines indicating best-fit linear regression (panels A and B). Alternatively, interaction analysis for age ($n = 62$ for < 70 yr + $n = 28$ for ≥ 70 yr) and maximal ascending aortic diameter ($n = 32$ for < 40 mm + $n = 58$ for ≥ 40 mm) on (**C**) peak and (**D**) plateau delamination pressures are shown for individual specimens, $n = 90$ ($n = 24$ for diameter < 40 mm and age < 70 yr, $n = 38$ for diameter ≥ 40 mm and age < 70 yr, $n = 8$ for diameter < 40 mm and age ≥ 70 yr, and $n = 20$ for diameter ≥ 40 mm and age ≥ 70 yr), two-way ANOVA with Tukey's multiple comparison test (panels C and D), ns: not significant, *$p < 0.05$, **$p < 0.01$, ***$p < 0.001$, ****$p < 0.0001$. There was no significant interaction between the independent variables of age and aortic diameter for the dependent variables of peak ($p = 0.25$) or plateau ($p = 0.11$) pressures, suggesting additive effects.

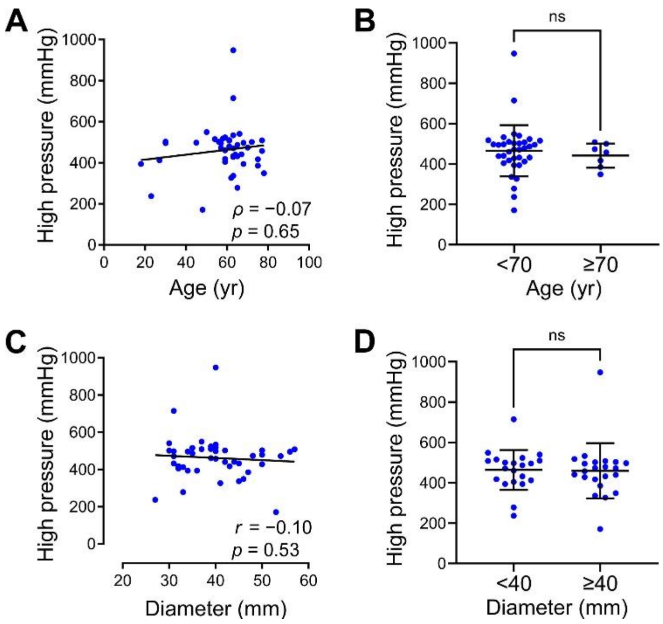

**Supplemental Figure 5: Age and aortic diameter do not correlate with injection pressures for medial extravasation.** Terminal high pressures required for medial extravasation at low flow rate in ascending aorta specimens from organ donors and surgical patients were compared to age and maximal ascending aortic diameter. (**A**) Age did not correlate with extravasation pressures, and (**B**) aortas from subjects younger or older than 70 years extravasated at similar pressures. (**C**) Aortic diameter did not correlate with extravasation pressures, and (**D**) aortas with diameters smaller or larger than 40 mm extravasated at similar decreased pressures. Data for individual specimens are shown, $n = 43$ ($n = 36$ for age < 70 yr, $n = 7$ for age ≥ 70 yr, $n = 21$ for diameter < 40 mm, and $n = 22$ for diameter ≥ 40 mm), ns: not significant, $\rho$ correlation coefficient and $p$-value by Spearman correlation with lines indicating best-fit linear regression (panel A), Mann-Whitney test (panel B), $r$ correlation coefficient and $p$-value by Pearson correlation with lines indicating best-fit linear regression (panel C), and unpaired t-test (panel D).

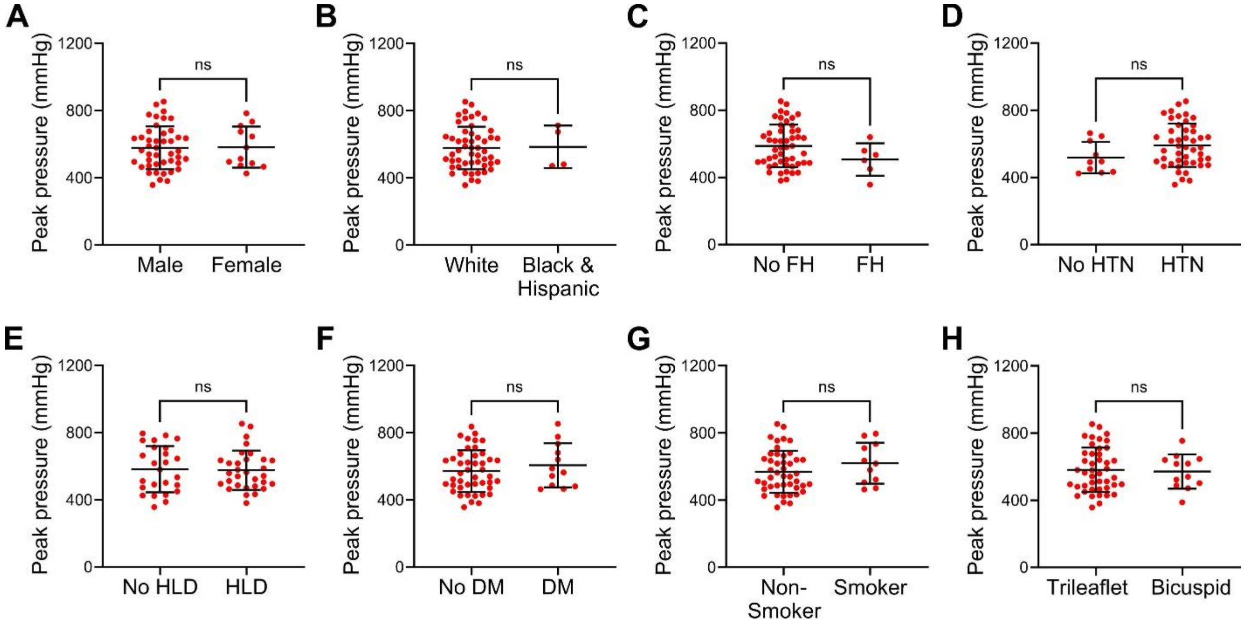

**Supplemental Figure 6: Demographic and clinical determinants of peak pressures required for medial delamination by subjects.** Peak pressures required to initiate medial delamination at high flow rate in ascending aortas from organ donors and surgical patients were compared by demographic and clinical factors with categorical variables. There was no difference in peak pressures among (**A**) male vs. female subjects, and (**B**) white vs. black and Hispanic subjects. There was also no difference in peak pressures among subjects with (**C**) family history (FH) of thoracic aortic aneurysm or dissection, (**D**) hypertension (HTN), (**E**) hyperlipidemia (HLD), (**F**) diabetes mellitus (DM), (**G**) smoker status, and (**H**) presence of bicuspid aortic valve. Data for individual subjects are shown, $n$ = 56 ($n$ = 44 male + $n$ = 12 female, $n$ = 52 white + $n$ = 4 black and Hispanic, $n$ = 50 no FH + $n$ = 6 FH, $n$ = 10 no HTN + $n$ = 46 HTN, $n$ = 25 no HLD + $n$ = 31 HLD, $n$ = 44 no DM + $n$ = 12 DM, $n$ = 45 non-smoker + $n$ = 11 smoker, and $n$ = 44 trileaflet + $n$ = 12 bicuspid), ns: not significant, unpaired t-test.

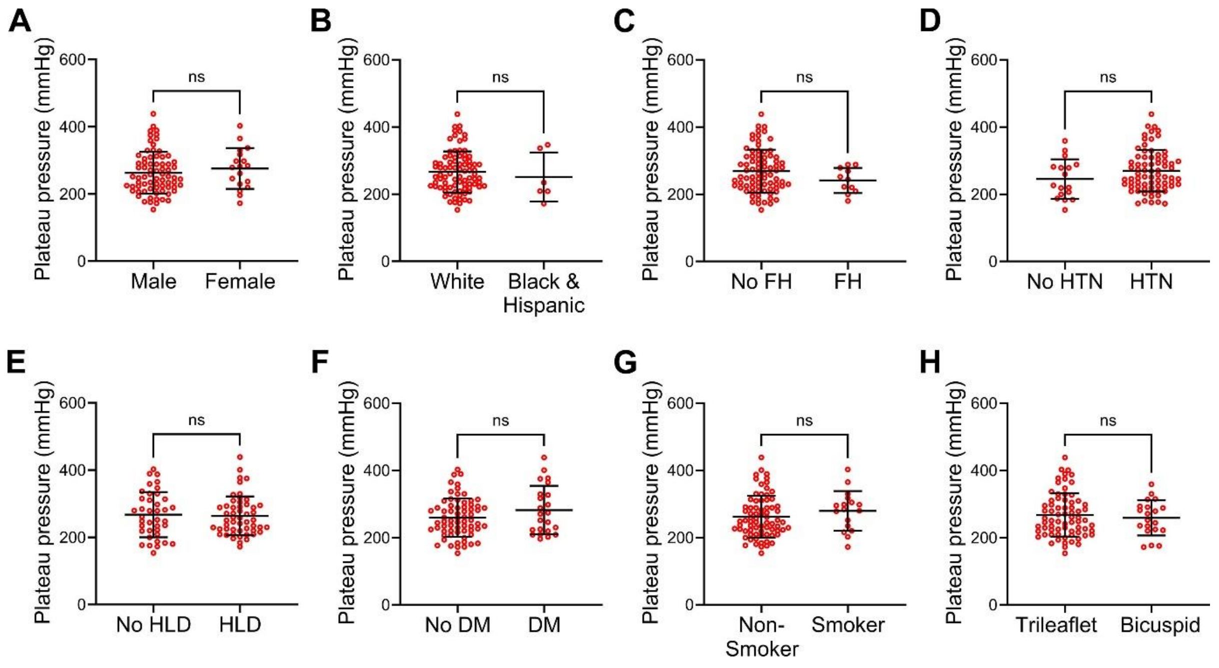

**Supplemental Figure 7: Demographic and clinical determinants of plateau pressures required for medial delamination by specimens.** Plateau pressures required to propagate medial delamination at high flow rate in ascending aortas from organ donors and thoracic aortic aneurysm patients were compared by demographic and relevant clinical variables. (**A**) Sex, (**B**) race/ethnicity, (**C**) family history (FH) of thoracic aortic aneurysm or dissection, (**D**) hypertension (HTN), (**E**) hyperlipidemia (HLD), (**F**) diabetes mellitus (DM), (**G**) smoker status, and (**H**) presence of bicuspid aortic valve had no impact on plateau pressures. Data for individual specimens are shown, $n = 90$ ($n = 72$ male + $n = 18$ female, $n = 84$ white + $n = 6$ black and Hispanic, $n = 80$ no FH + $n = 10$ FH, $n = 17$ no HTN + $n = 73$ HTN, $n = 38$ no HLD + $n = 52$ HLD, $n = 67$ no DM + $n = 23$ DM, $n = 73$ non-smoker + $n = 17$ smoker, and $n = 71$ trileaflet + $n = 19$ bicuspid), ns: not significant, unpaired t-test.

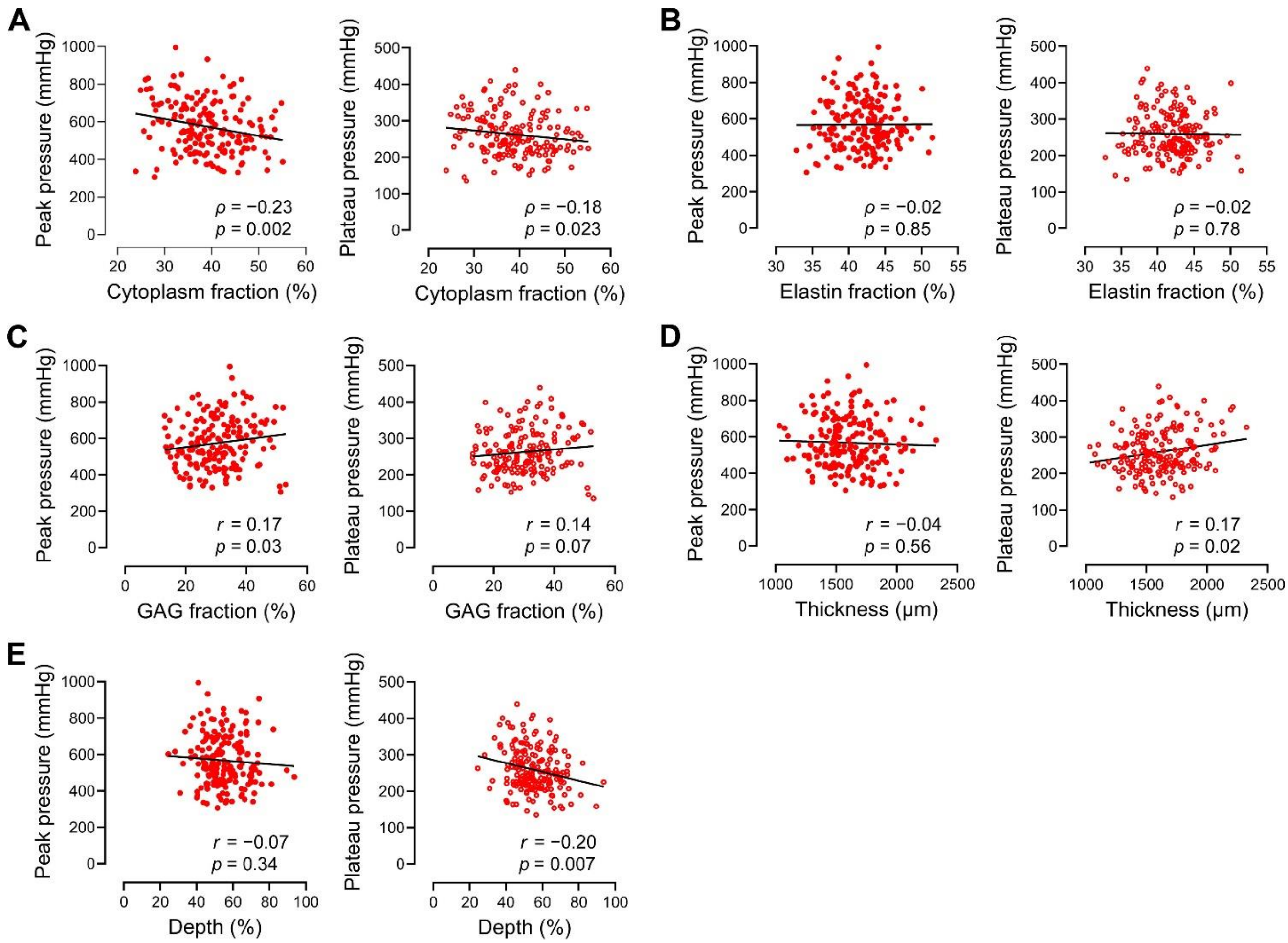

**Supplemental Figure 8: Correlations of medial components and dimensions with delamination pressures.** Histological measurements on Movat's pentachrome stains were compared to peak and plateau pressures required to initiate and propagate medial delamination at high flow rate. (**A**) Increased cytoplasm fraction in the aortic media correlated with decreased peak and plateau pressures. (**B**) Elastin fraction had no impact on delamination pressures. (**C**) Increased glycosaminoglycan (GAG) fraction weakly correlated with increased peak pressure; there was not a significant association with plateau pressure. (**D**) Increased medial thickness weakly correlated with increased plateau but not peak pressure. (**E**) The relative depth of the major delamination plane (distance of intima to major delamination plane relative to total medial thickness) had a negative association with plateau but not peak pressure. Data for individual injections are shown, $n$ = 173–184 depending on suitable quality of histological stains for image quantification, $\rho$ correlation coefficient and $p$-value by Spearman rank correlation with lines indicating best-fit linear regression (panels A, B), and $r$ correlation coefficient and $p$-value by Pearson correlation with lines indicating best-fit linear regression (panels C–E).

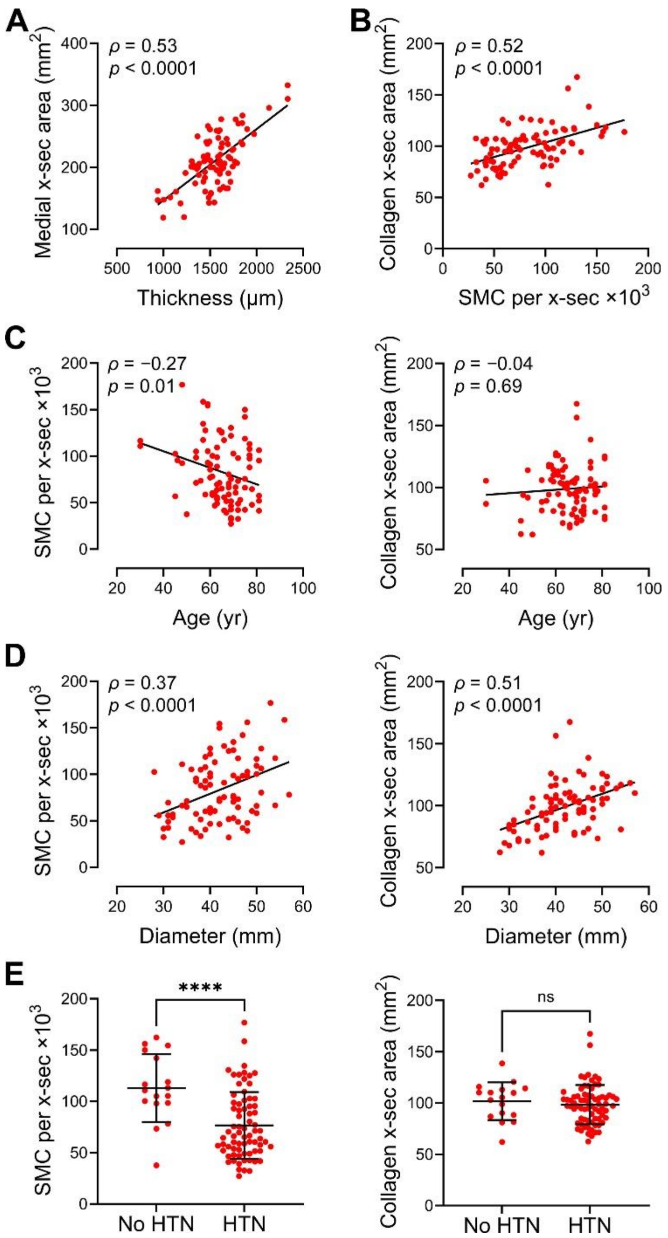

**Supplemental Figure 9: Relationship of clinical factors and cross-sectional amounts of medial components.** Total number of cells and collagen content were calculated per cross-section of aorta. There were strong correlations between (**A**) medial thickness and medial cross-sectional (x-sec) area and (**B**) collagen content per cross-section and number of smooth muscle cells (SMC) per cross-section. (**C**) Ageing correlated with decreased number of cells but not collagen content per cross-section. (**D**) Aortic dilatation correlated with increased smooth muscle cells and collagen per cross-section. (**E**) Hypertension (HTN) associated with decreased number of cells but not collagen content per cross-section. Data for individual specimens are shown, $n$ = 90, $\rho$ correlation coefficient and $p$-value by Spearman correlation with lines indicating best-fit linear regression (panels A–D), and unpaired t-test (panel E).

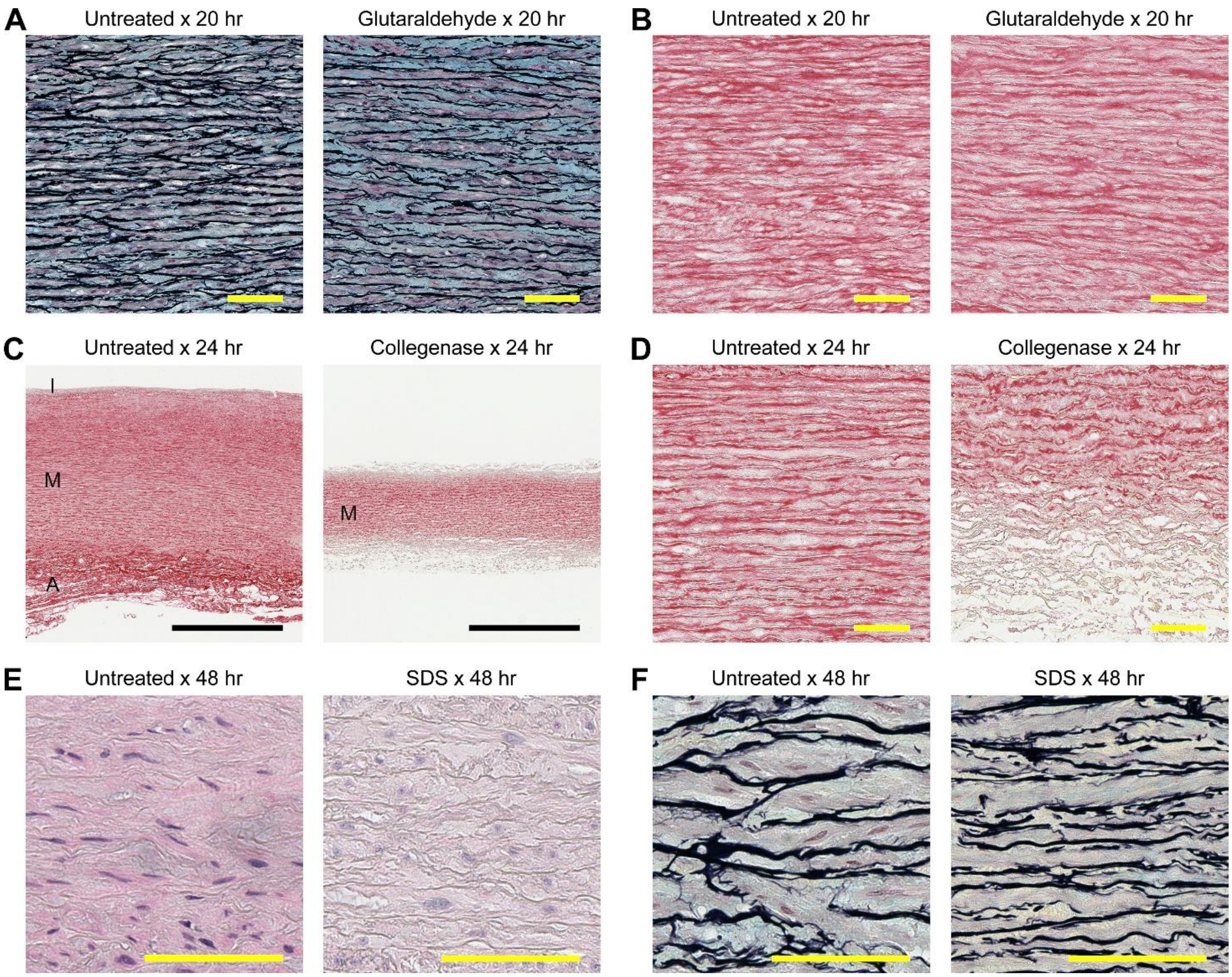

**Supplemental Figure 10: Pretreatment of aortic specimens to increase protein cross-linking, decrease collagen content, or decrease cell numbers.** Ascending aortic specimens were pretreated with various agents with adjacent tissue pieces remaining untreated as controls prior to intramural fluid injection. (**A**) Movat and (**B**) sirius red stains of untreated or glutaraldehyde-treated aortic specimens after 20 hr showing no overt changes to medial cells and extracellular matrix from fixation. (**C**) Sirius red stains at low magnification or (**D**) high magnification of untreated or collagenase-treated aortic specimens after 24 hr showing substantial thinning of media (M) from both intima (I) and adventitia (A) surfaces with heterogeneous loss of collagen most pronounced in the remaining superficial laminae. (**E**) Hematoxylin and eosin and (**F**) Movat stains of untreated or SDS-treated aortic specimens after 48 hr showing loss of cellular and nuclear features but not of extracellular matrix from detergent. Scale bars represent 100 μm (panels A, B, D, E, F) or 1 mm (panel C). Results are representative of paired specimens from non-dilated aortas of 6 subjects.
